# HP1-driven phase separation recapitulates the thermodynamics and kinetics of heterochromatin condensate formation

**DOI:** 10.1101/2022.07.11.499635

**Authors:** Maxime M.C. Tortora, Lucy D. Brennan, Gary Karpen, Daniel Jost

## Abstract

The spatial segregation of pericentromeric heterochromatin (PCH) into distinct, membrane-less nuclear compartments involves the binding of Heterochromatin Protein 1 (HP1) to H3K9me2/3-rich genomic regions. While HP1 exhibits liquid-liquid phase separation properties *in vitro*, its mechanistic impact on the structure and dynamics of PCH condensate formation *in vivo* remains largely unresolved. Here, using biophysical modeling, we systematically investigate the mutual coupling between self-interacting HP1-like molecules and the chromatin polymer. We reveal that the specific affinity of HP1 for H3K9me2/3 loci facilitates coacervation *in nucleo*, and promotes the formation of stable PCH condensates at HP1 levels far below the concentration required to observe phase separation in purified protein assays *in vitro*. These heterotypic HP1-chromatin interactions give rise to a strong dependence of the nucleoplasmic HP1 density on HP1-H3K9me2/3 stoichiometry, consistent with the thermodynamics of multicomponent phase separation. The dynamical crosstalk between HP1 and the viscoelastic chromatin scaffold also leads to anomalously-slow equilibration kinetics, which strongly depend on the genomic distribution of H3K9me2/3 domains, and result in the coexistence of multiple long-lived, microphase-separated PCH compartments. The morphology of these complex coacervates is further found to be governed by the dynamic establishment of the underlying H3K9me2/3 landscape, which may drive their increasingly abnormal, aspherical shapes during cell development. These findings compare favorably to 4D microscopy measurements of HP1 condensates that we perform in live *Drosophila* embryos, and suggest a general quantitative model of PCH formation based on the interplay between HP1-based phase separation and chromatin polymer mechanics.

**SIGNIFICANCE STATEMENT:** The compartmentalization of pericentromeric heterochromatin (PCH), the highly-repetitive part of the genome, into membrane-less organelles enriched in HP1 proteins, is critical to both genetic stability and cell fate determination. While HP1 can self-organize into liquid-like condensates *in vitro*, the roles of HP1 and the polymer chromatin in forming 3D PCH domains *in vivo* are still unclear. Using molecular simulations, we show that key kinetic and thermodynamic features of PCH condensates are consistent with a phase-separation mode of organization driven by the genomic distribution of methylated domains and HP1 self-attraction and affinity for heterochromatin. Our predictions are corroborated by live-microscopy performed during early fly embryogenesis, suggesting that a strong crosstalk between HP1-based phase separation and chromosome mechanics drive PCH condensate formation.

## INTRODUCTION

The regulation of genome function in eukaryotes is a highly complex biological process, which typically involves the combined contribution of hundreds of intra-nuclear proteins, DNA and RNA structures (1). The precise and coordinated recruitment of regulatory molecules at specific genomic or molecular targets often gives rise to spatially- and biochemically-distinct compartments, which play essential roles in a variety of cellular activities including DNA repair (2), transcription (3, 4), replication (5) and epigenetic regulation (6). These membrane-less condensates act as both organizational hubs and localized crucibles for the catalysis of multiple biochemical reactions involved in the translation and maintenance of genetic information (7). In recent years, biophysical characterizations of some condensates (e.g., P-granules) suggested that they behave as phase-separated liquids with respect to structure (e.g., round shapes, concentration-dependent assembly) and dynamics (e.g., rapid exchange of molecules with the environment, ability to fuse when in contact) (8). Therefore, the physical mechanism of liquid-liquid phase separation (LLPS) has been increasingly proposed as a key paradigm to explain their formation based on inherent interactions among solvent, proteins, nucleic acids (RNA and DNA), or other biomolecules (7, 9, 10). However, in some other cases, the difficulty of quantitatively and unequivocally distinguishing LLPS from alternative self-organization processes *in vivo* has generated disputes about its biological relevance (11–13).

A particularly notable debate has focused on the case of pericentromeric heterochromatin (PCH), which forms large, distinct nuclear compartments required for chromosome folding, mitotic segregation, and transcriptional silencing of transposons and genes (14). Pericentromeres are biochemically defined by genomic regions enriched in repeated DNA sequences, as well as in histone H3 lysine 9 di- or tri-methylation marks (H3K9me2/3) and their epigenetic “reader” protein, Heterochromatin Protein 1a/α (HP1a/α) (15). Careful genetic and biochemical studies have teased apart a hierarchy of interactions that contribute to the formation and maintenance of pericentromeric domains.

The first level of interactions involves the dimerization of HP1a/α molecules via their *chromoshadow* domains (CSDs) (16) and the direct binding of HP1a/α dimers to H3K9me2/3 (hereafter referred to as “methylated chromatin”) mediated by their *chromodomains* (CDs) (17). The high-affinity dimerization interactions have led the field to consider the HP1 dimer to be the functional form across homologs (18). The ability of HP1 molecules to form stable dimers also underlies a well-supported structural model for pericentromeres, whereby adjacent methylated nucleosomes may be transiently bridged by HP1 dimers (19, 20) – thus sequestering the underlying chromatin fiber into distinct, compact spatial compartments (henceforth referred to simply as “PCH condensates”).

The second level of interactions concerns a complex set of low-affinity interactions driven primarily by the multiple intrinsically-disordered regions (IDRs) within the HP1a/α dimer, which in turn modulate the interaction strengths of both HP1 structured domains (CSD & CD) and the interactions with nucleic acids and other proteins (21, 22). Extensive investigations in mammals and Drosophila – based on HP1α and HP1a, respectively – have demonstrated that such interactions may drive the ability of HP1 to spontaneously form distinct, liquid-like condensates *in vitro* (23–25).

All of these interactions are further affected by ionic conditions, nucleic acid content and accessibility, post-translational modifications such as HP1α phosphorylation, and HP1 protein-partner binding (23, 25) suggesting a multiplicity of possible mechanisms for regulating the *in vivo* segregation of methylated chromatin into PCH compartments.

Currently, two main classes of mechanisms have been hypothesized (12) (Fig.1a): a “LLPS-like” mode of organization where PCH compartments result mainly from the capacity of HP1-like architectural proteins to self-interact and to form liquid-like droplets within which H3K9me2/3-rich regions may colocalize; and a “bridging-like” process where condensates emerge solely from the capacity of HP1 to carry multivalent heterotypic bonds with methylated chromatin, and hence to stabilize direct, transient “bridges” between distant loci. The former is backed up by *in vivo* observations in mammals and Drosophila revealing a highly-dynamic exchange of HP1 molecules within PCH and in the nucleoplasm (26), and the rapid kinetics of PCH assembly and disassembly during the cell cycle (25, 27). While the latter is supported by *in vivo* studies in mammalian cell lines suggesting that the structural properties of HP1 condensates deviate from those observed *in vitro* in LLPS-driven solutions of purified proteins and resemble more that of polymer globules with size and compaction only weakly dependent on HP1 concentration (28). However, this assumed dichotomy between bridging- and LLPS-driven compartmentalization does not account for the complex physics of intra-cellular multicomponent phase separation (29), which may significantly differ from that of simple single-component LLPS phase separation within the crowded nuclear environment (11, 29, 30).

**Figure 1:**
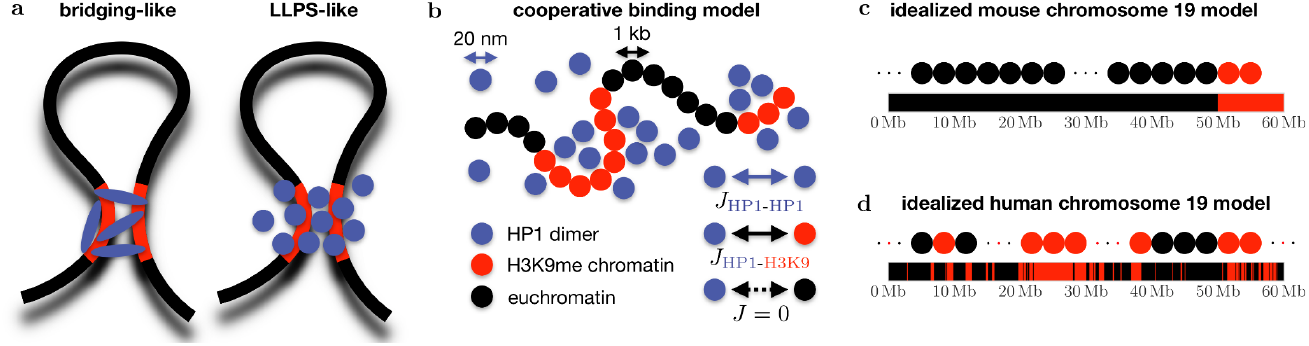
Cooperative binding model: chromosome folding driven by self-interacting proteins. (a) Two possible pathways for the 3D compartmentalization of genomic loci sharing the same epigenomic content: (left) a bridging-like process where multivalent chromatin-binding proteins directly link two distant regions; (right) a LLPS-like mechanism where self-interacting binders form condensates within which a specific type of chromatin may colocalize. (b) Sketch of the cooperative binding model: diffusible molecules (HP1 dimers) may self-interact (*J*_*HP1*− *HP*1_) and have an affinity (*J*_*HP*1−*H*3*K*9_) for specific regions (H3K9me2/3) of a polymer (chromatin). (c,d) Representations of the idealized polymer models of a telocentric mouse chromosome 19 (c) and human chromosome 19 (d).

Here, we aim to reconcile these *a priori* conflicting *in vitro* and *in vivo* observations of HP1-driven phase separation through the lens of a generic first-principle model of endogenous chromatin-protein assemblies. Actually, the vast majority of theoretical studies to date have investigated the formation of PCH compartments (and of the 3D genome in general) using “bridging-like” models (31–37) and the generic physics of the LLPS-like mode of organization is poorly characterized (30). Our minimal biophysical description enables us to comprehensively explore the basic physics of heterogeneous, multicomponent coacervates composed by chromatin and architectural proteins, and is contextualized to address the differential roles of HP1-HP1 and HP1-H3K9me2/3 interactions in PCH assembly. In particular, we demonstrate that the key kinetic and thermodynamic features of PCH condensate formation are compatible with a mode of organization in which chromosome structure and dynamics are driven by the coupling between fast-diffusing HP1 proteins, H3K9me2/3 patterning along the chromosome, and the constrained chromatin fiber. Our theoretical predictions are corroborated by *in vivo* 4D microscopy measurements of HP1a condensate formation in early Drosophila embryos, and may provide a versatile framework to interpret the endogenous behavior of a wide range of biomolecular condensates.

## RESULTS

### A minimal model for chromosome folding driven by self-interacting proteins

To investigate the potential interplay between the self-affinity of architectural chromatin-binding proteins such as HP1 and the large-scale organization of chromosomes, we developed a generic biophysical framework (Fig.1b) – termed the *cooperative binding model* – which accounts for the coupled dynamics of the chromatin polymer and self-interacting diffusible particles (see Materials & Methods for details). Briefly, we represent chromatin as a self-avoiding, semi-flexible chain (38) and describe protein binders via the lattice gas model, which corresponds to an efficient molecular-level description of phase separation (39). The spatio-temporal evolution of the system is driven by standard polymer properties (backbone connectivity, excluded volume and bending rigidity), in conjunction with both homotypic, HP1-HP1 interactions and a heterotypic affinity of HP1 for H3K9me2/3 monomer sites along the chromatin chain. In this context, diffusible particles represent HP1 dimers with a typical hydrodynamic radius of 20 nm, and each target monomer (referred to as a *locus*) corresponds to a contiguous chromatin region of length ∼1 kbp, in which all constituent histones are assumed to bear the H3K9me2/3 post-translational mark (see Materials & Methods).

### The lattice-gas model recapitulates *in vitro* features of HP1-based phase separation

To assess the ability of the model to reproduce some of the main kinetic and equilibrium properties of classical, single-component phase separation, we first performed simulations involving pure HP1 dimers in the absence of the chromatin scaffold (Fig.2a). For this purpose, we systematically varied the total HP1 concentration *ρ*_*HP*1_ and self-affinity *J*_*HP*1−*HP*1_ between HP1-dimers, and monitored both the time evolution and resulting steady state of the system, starting from a well-mixed random initial state. We find that, beyond a critical strength of self-affinity (*J*_*crit*_ ≃ 1 kJ/mol), the value 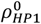 of HP1 concentration above which condensates form in our simulations is a steep function of *J*_*HP*1−*HP*1_, and decreases monotonously from 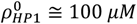 to 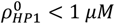 upon increasing *J*_*HP*1−*HP*1_ in the range [1 − 3] *kJ*/*mol* (Fig.2b, black curve). We observe that increasing the total HP1 content *ρ*_*HP*1_ induces a swelling of the condensate (Fig.2a), while the HP1 level 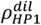 outside the focus similarly remains fixed (Fig.2c). This *concentration buffering* effect signifies that, in the presence of condensates, the concentration of the dilute phase is independent of the total HP1 content of the system (11, 29, 40).

**Figure 2:**
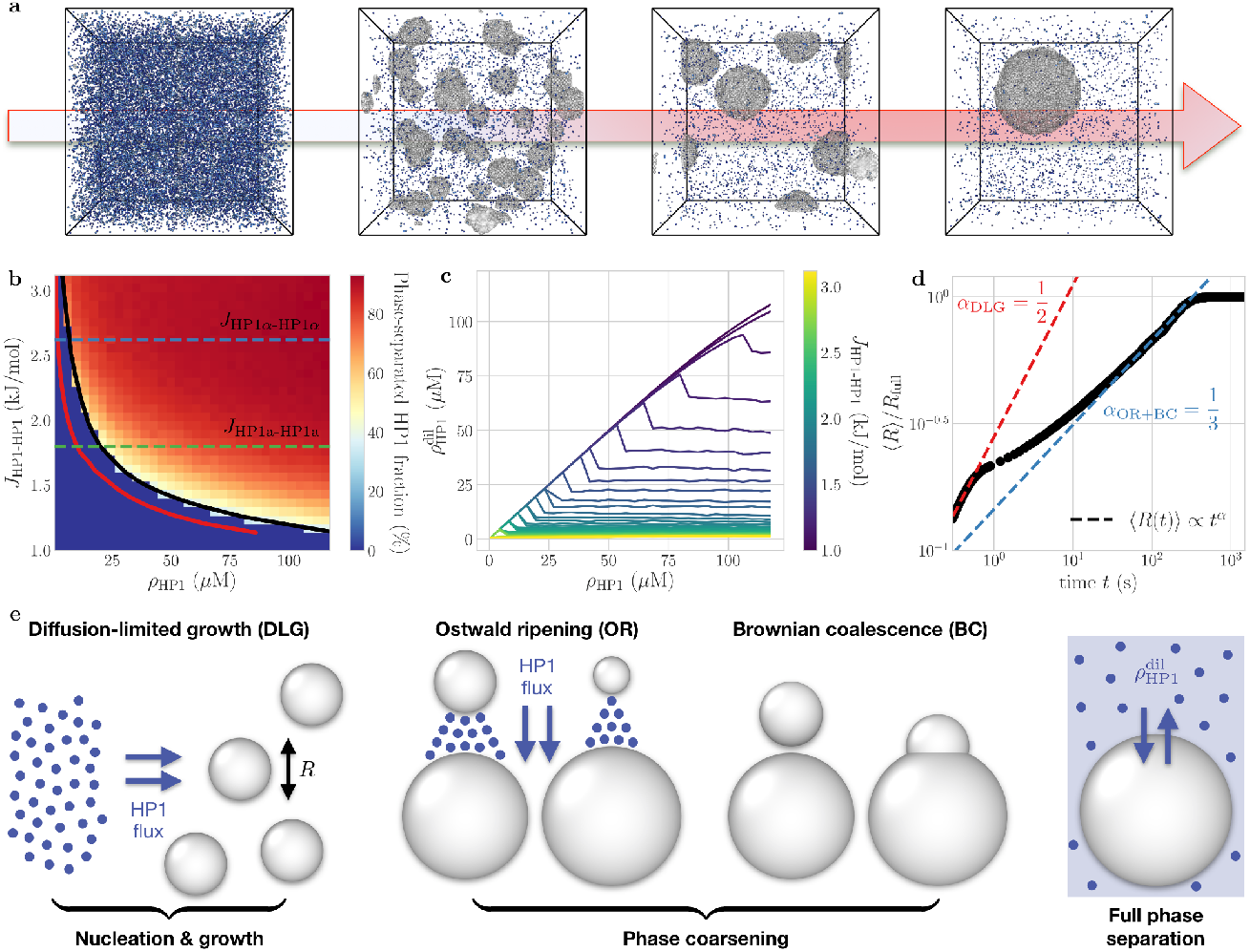
Simulations recapitulates key features of single-component phase separation. (a) Typical kinetic pathway of the simulations, starting from a well-mixed, homogeneous initial state, for a set of parameters *ρ*_*HP*1_= 8 *μM, J*_*HP*1−*HP*1_ = 2.6 kJ/mol located within the two-phase coexistence region (see Movie S1). (b) Phase diagram of pure HP1 system showing the fraction of HP1 in condensates as a function of HP1 density *ρ*_*HP*1_ and homotypic affinity *J*_*HP*1−*HP*1_. Red area: two-phase coexistence region in our simulation. Red line: HP1 concentration in the dilute phase (see (c)). Dashed lines: affinities inferred for HP1α (blue) and HP1a (green), respectively (see text). (c) Background HP1 density 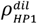 in the dilute phase as a function of total HP1 level *ρ*_*HP*1_. Each line corresponds to a fixed value of *J*_*HP*1−*HP*1_. (d) Kinetics of the average condensate radius normalized by its equilibrium value *R*_*full*_. Simulation parameters are as in (a). (e) Standard phase separation equilibration mechanisms reproduced by the model: from rapid, diffusion-limited growth at early times to slower, coarsening-dominated behaviors at later stages.

Then, we monitored the kinetics of HP1 condensates predicted by the model close to the stability line (Fig.2b, black curve). It displays a marked transition from an initiation and growth stage at short times (*t* ≲ 1 *s*) to a coarsening behavior at longer times (*t* ≳ 1 *s*), in agreement with classical theories of phase separation dynamics (8) (Fig.2a). The initial stage is characterized by a diffusion-limited growth (DLG) process, associated with a rapid expansion of the mean condensate size ⟨*R*(*t*)⟩ ∝ *t*^1/2^ (Fig.2d,e) (8), constrained by the diffusion rate of individual proteins within the bulk. At later time points, the coarsening regime describes the subsequent growth of larger HP1 droplets at the expense of the smaller foci, either via local collision events driven by the stochastic diffusion of proximal foci – referred to as Brownian coalescence (BC) – or via non-local, transient protein “evaporation” and recondensation, known as Ostwald ripening (OR) (Fig.2e) (8). Both mechanisms are associated with a slower growth exponent ⟨*R*(*t*)⟩ ∝ *t*^1/3^ (Fig.2d) (8) and can be observed in our simulations (Fig.2a), although OR generally constitutes the dominant coarsening process in this simple lattice-gas framework (see Materials & Methods).

Altogether, these results evidence the capacity of the model to recapitulate salient features of classical single-component phase-separation, both in- and out-of-equilibrium. We finally used such analysis to parametrize HP1-HP1 interactions based on previous *in vitro* investigations for purified HP1α and HP1a proteins using turbidity assays in physiological buffer conditions (23, 25, 28). Identifying 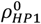 as the HP1 density at the onset of turbidity (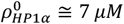 for mouse HP1α and 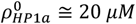 for Drosophila HP1a, see SI Methods), we find *J*_*HP*1*α*−*HP*1*α*_ ≅ 2.6 kJ/mol and *J*_*HP*1*a*−*HP*1*a*_ ≅ 1.8 kJ/mol, respectively (Fig.2b). These values are consistent with the weak, reversible inter-protein interactions typically characterizing LLPS (10, 41).

### HP1-H3K9me2/3 affinity impacts the thermodynamics of HP1 condensates *in nucleo*

Having in hand a minimal model of homotypic HP1 interactions, we investigated how the formation of HP1 condensates *in vivo* may be impacted by the heterotypic affinity between HP1 dimers and H3K9me2/3-enriched genomic loci (Fig.1b, 3a), and how it may in turn impact the structure and dynamics of PCH compartments (Fig.3b-g, see also Fig.S4). To that end, we consider as a first test case a simplified representation of mouse chromosome 19 (60 *Mbp*) featuring a unique, 10 *Mbp*-long H3K9me2/3 telocentric domain (Fig.1c) in the presence of HP1α. We report in Fig.3b that the addition of this methylated chromatin domain considerably decreases the concentration above which HP1 foci form in simulations. Indeed, HP1 condensates may now be observed at intra-nuclear HP1 levels far lower than the density 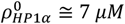 required *in vitro* to phase-separate purified HP1α (Figs.2b,3b). This effect is found to be increasingly pronounced upon raising *J*_*HP*1−*H*3*K*9_ in the range [0.1 − 0.6] *kJ*/*mol*, which corresponds to the typical magnitude of HP1-H3K9 binding energy suggested by previous computational studies (32). Coincidentally, we find that the radius *R*_0_ of the smallest stable HP1 foci in our simulations is also significantly reduced upon increasing *J*_*HP*1−*H*3*K*9_ (Fig.3d). Both observations can be attributed to the binding of HP1 onto the methylated chromatin substrate, which promotes phase separation by facilitating focus nucleation and compensates for the higher interfacial energy costs (Fig.S1) associated with smaller droplets.

**Figure 3:**
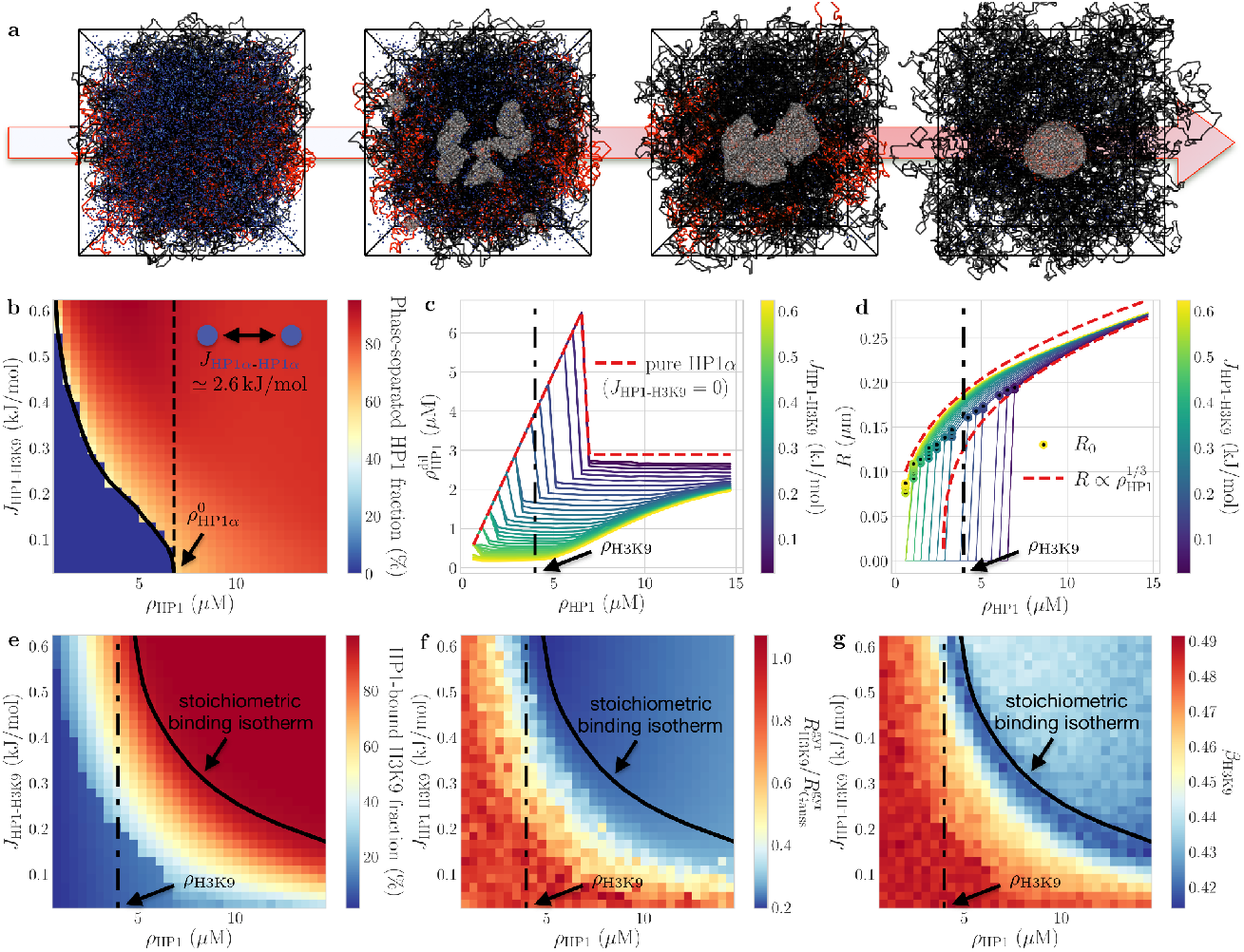
HP1-H3K9me2/3 interactions impact condensate thermodynamics. Case of an idealized telocentric mouse chromosome (Fig.1c). (a) Typical kinetic pathway of the simulations (*ρ*_*HP*1_ = 8 *μM, J*_*HP*1−*H*3*K*9_ = 0.5 kJ/mol, *J*_*HP*1−*HP*1_ = 2.6 kJ/mol) (see Movie S2). (b) Phase diagram of HP1α in presence of chromatin (fraction of HP1 in condensates) as a function of *ρ*_*HP*1_. Each line corresponds to a fixed value of *J*_*HP*1−*H*3*K*9_. Red area: two-phase coexistence region. Dashed line: *in vitro* threshold concentration 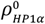. (c) Background HP1 level 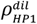 in the dilute (nucleoplasmic) phase as a function of total HP1 density *ρ*_*HP*1_ at various fixed *J*_*HP*1−*H*3*K*9_. Red dashed line: values computed for pure HP1α dimers (Fig.2c). (d) Same as (e) but for the average size *R* of HP1-dense foci. Dashed lines: growth behavior expected for a concentration-buffered system in the limits of high and low *J*_*HP*1−*H*3*K*9_ (see SI Methods). *R*_0_ denotes the radius of the smallest stable focus observed for each *J*_*HP*1−*H*3*K*9_ value. (e) Fraction of methylated monomers bound to at least one HP1 dimer. Black full line: minimal (*J*_*HP*1−*H*3*K*9_, *ρ*_*HP*1_) values at which this fraction reaches 100% (stoichiometric binding isotherm). (f) Radius of gyration 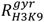 of the HP1 domain, normalized by the expected value 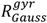 for a non-interacting, Gaussian coil. (g) Diffusion exponent *β*_*H*3*K*9_ of methylated loci 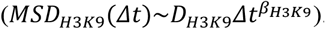. The black dash-dotted lines in (c-g) represent *ρ*_*H*3*K*9_, the level of nuclear methylated chromatin imposed in the simulations.

Furthermore, in contrast to the pure HP1 system, the potential concentration buffering behavior is now dependent on the system composition (Fig.3c). For moderate-to-high affinity strengths *J*_*HP*1−*H*3*K*9_, we observe that the HP1 density 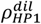 in the dilute, nucleoplasmic phase remains almost constant for low HP1 levels, then increases with *ρ*_*HP*1_– and only approaches its *in vitro* plateau value at very high concentrations. To interpret this stoichiometric effect, we define the *stoichiometric binding isotherm* as the minimal (*J*_*HP*1−*H*3*K*9_, *ρ*_*HP*1_) values at which all the methylated loci are in contact with at least one HP1 dimer (Fig.3e, black line). Below the isothermal value, the system is in a regime of droplet growth driven by HP1-H3K9 binding events and most of the HP1 dimers are wetting H3K9me2/3 sites, the population of free dimers being negligible 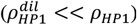 (Fig.3c). Beyond this point, additional dimers chiefly diffuse into the nucleoplasmic background phase, as further swelling of the droplets does not incur any gains in HP1-H3K9 contacts. Thus, one generally obtains a sublinear growth of the focus volume *V* with increasing *ρ*_*HP*1_ (Fig.3d), and only recovers the linear growth regime of standard concentration buffering in the limit of wide HP1 excess (*ρ*_*HP*1_ ≫ *ρ*_*H*3*K*9_) – in which HP1-H3K9me2/3 coupling becomes statistically irrelevant, and is superseded by HP1-HP1 homotypic interactions (Figs.3c,d).

The formation of HP1 condensates is further found to drive the partial or total collapse of the encapsulated methylated chromatin (Fig.3f). This structural transition starts when roughly 20-30% of the H3K9me2/3 loci are covered by HP1 dimers, and is also associated with a dynamical crossover. At this point, the methylated chromatin diffusion constant is reduced by roughly fivefold (Fig.S3f) and the diffusion exponent goes from 0.5 – consistent with the viscoelastic dynamics of a dilute, unconstrained polymer – down to ∼0.4, characterizing the slower motion of a dense, globular chain (Fig.3g) (42). Interestingly, the stoichiometric binding region coincides with a maximal impact on the compaction and mobility of the methylated chromatin region (Figs.3f,g and S2h), which results from the full encapsulation of all H3K9me2/3 loci within a HP1 focus of minimal volume, and thus leads to the maximal degree of PCH packaging.

Together, these conclusions demonstrate that the ability of HP1 dimers to self-interact, coupled with the specific affinity of HP1 for methylated chromatin, mediates the formation of condensed PCH compartments at endogenous levels far below the concentrations required to observe HP1 condensation in purified *in vitro* assays. This result also highlights the limitations of intra-cellular perturbation experiments based on ectopic protein over- or under-expression in investigations of *in vivo* phase separation (28), whose interpretation in terms of concentration buffering generally requires a careful quantitative analysis of the levels of all the interaction partners involved (29).

### H3K9me2/3 distribution and chromosome mechanics regulate phase separation kinetics

The coupled evolution dynamics of the HP1-chromatin system studied in Fig.3 irreversibly leads to rapid relaxation towards a fully phase-separated state, characterized by the complete segregation of methylated chromatin into a single, large focus encapsulated within a unique HP1 liquid droplet (Fig.3a, right). The phase separation kinetics predicted by the model in this case reveal a quick equilibration of the mean focus size over a typical timescale *t* ≲ 10 *s* with a growth exponent ⟨*R*(*t*)⟩ ∝ *t*^1/2^, consistent with nucleation-driven DLG (Fig.4e). We attribute this effect to the cooperative binding of HP1 onto the contiguous methylated chromatin domain, which leads to the strong co-localization of droplet nucleation sites within the same, H3K9me2/3-dense nuclear region (Fig.3a, left) – and thus results in rapid growth independent of the slower OR and BC coarsening processes.

**Figure 4:**
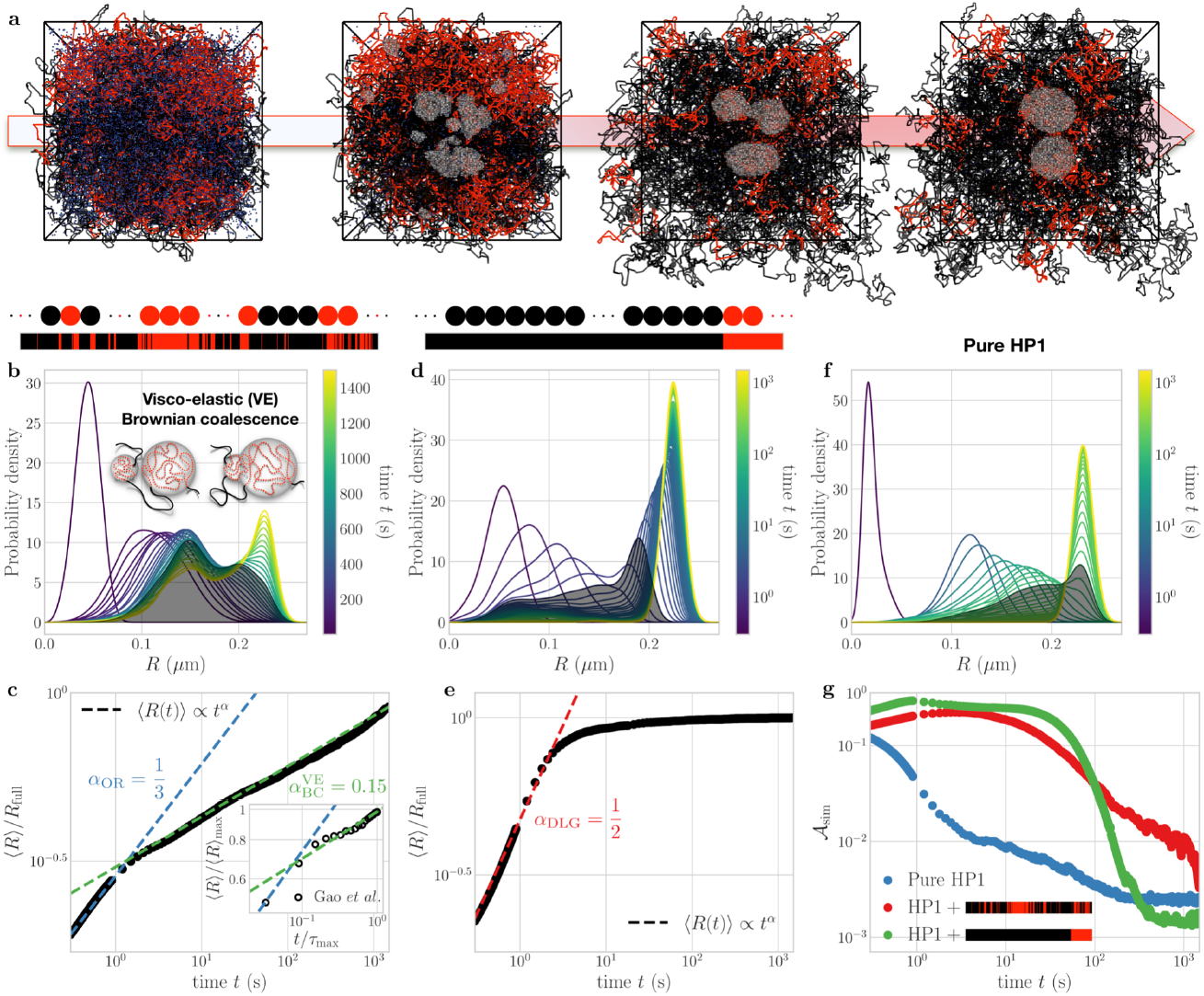
H3K9me2/3 distribution and chromatin mechanics govern phase separation kinetics. (a) Typical kinetic pathway of the simulations in the case of idealized polymer model of human chromosome 19 (see Movie S3). (b) Time evolution of distributions of HP1 droplet size *R* in the human chromosome 19 system. Inset: Illustration of the viscoelastic (VE) Brownian coalescence process. (c) Associated evolution kinetics of the mean droplet radius ⟨*R*⟩ normalized by its equilibrium value *R*_*full*_. Inset: Experimental data from Ref. (43), obtained from the mean fluorescence intensity of engineered HP1 puncta (see SI Methods). (d,e) Same as (b,c) in the case of an idealized telocentric mouse chromosome. (f) Same as (b) for pure HP1α dimers (c.f. Fig.2). (i) Time evolution of the focus anisotropy *A*_*sim*_, such that *A*_*sim*_ = 0 for ideal spheres and *A*_*sim*_ → 1 for elongated droplet shapes (see SI Methods). Simulation parameters as in Figs.2a & 3a.

While our first test case (Fig.1c) was representative of the formation of large chromocenters, it may be unable to describe the existing multiplicity of smaller, spatially-distinct HP1 compartments (44), observed during interphase in conventional mammalian nuclei (35). Such complex, heterogeneous domain patterns are traditionally interpreted in terms of *microphase separation* undergone by block copolymers in the presence of self-attractions between epigenomically-similar regions (45,91). To investigate whether such distinct compartments may be physically reconciled with a HP1-driven mechanism of heterochromatin folding, we introduced as a second test case a chromosome model featuring multiple methylated domains of disparate sizes, representative of chromosome 19 in human fibroblasts (Fig.1d, see Materials & Methods). We find that this heterogeneous distribution of H3K9me2/3 along the chromatin polymer leads to drastically-slower coarsening dynamics (Fig.4b,c). HP1 dimers first stochastically aggregate across multiple separate methylated regions. These competing nucleation sites then gradually mature into a number of distinct, spherical foci, which may remain metastable over timescales *t* ≫ 30 *min* well beyond the total simulation time. This partial phase coarsening process is characterized by the rapid, OR-like evaporation of smaller foci at short times (*t* < 1 *s*), associated with a mean growth exponent ⟨*R*(*t*)⟩ ∝ *t*^1/3^. Such events are however strongly suppressed at longer timescales (*t* > 2 *s*), over which focus growth is instead governed by an abnormally-slow growth exponent ⟨*R*(*t*)⟩ ∝ *t*^0.15^.

Similar anomalous coarsening dynamics have been recently reported for engineered intra-nuclear protein condensates, and were ascribed to the coupling of droplet motion with the viscoelastic (VE) diffusion kinetics of the underlying chromatin network (46–48). Indeed, the strong colocalization of H3K9me2/3 loci and HP1 foci (Fig.3a,e) imposes that condensate displacements correlate with the collective diffusion of the encapsulated methylated chromatin regions (Fig.4b, inset), and thus that individual HP1 foci experience a sub-diffusive motion with an exponent *β* similar to that of H3K9 domains (β ≅ 0.45, Fig.3g) (42). In this case, one may show (46) that the theoretical coarsening dynamics associated with diffusion of HP1 droplets simply scale as ⟨*R*(*t*)⟩ ∝ *t*^β/3^. The predicted scaling behavior 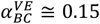 (Fig.4c) is thus consistent with such anomalously-slow, viscoelastic coalescence combined with an inhibition of OR, which we similarly attribute to the effective “trapping” of H3K9me2/3 domains within the HP1 condensates (49). Remarkably, a recent *in vivo* study in human cells based on CRISPR-engineered HP1 targeted to randomly-distributed ectopic chromatin binding sites (43) revealed that the assembly kinetics of HP1 foci at their multiple distinct target loci follow a similar transition from a *α* ≅ 1/3 short-time exponent to a slower growth regime with *α* ≅ 0.15 at longer times (Fig.4c, inset).

These findings suggest that anomalously-slow coarsening dynamics generically arise from the interplay between HP1-driven phase separation and chromosome mechanics in the presence of multiple, competing genomic binding regions. This conclusion is corroborated by an analysis of the time evolution of the distribution of droplet size (Fig.4b,d,f). In the case of pure HP1 (Fig.4f), for which OR dominates, distributions display a unimodal shape with a transient left-skewed long tail at intermediary times (1 *s* < *t* < 300 *s*), characteristic of the polydisperse population of small droplets undergoing continuous evaporation (8). In the presence of a single, large PCH genomic region (Fig.4d), population dynamics are qualitatively similar, which may be imputed to the rapid nucleation of HP1 foci onto the homogeneously-methylated chromatin substrate (50). Conversely, in the case of PCH combined with a highly-dispersed pattern of H3K9me2/3 marks (Fig.4b), we find that the tail is noticeably less pronounced, and that distributions instead display a transient bimodal shape at longer timescales. This behavior is consistent with the presence of metastable, separate HP1 foci that slowly merge into larger condensates via discrete coalescence events.

Thus, our results suggest that the microphase separation of HP1 and methylated regions into long-lived, coexisting nuclear compartments could be simply attributed to anomalously-slow equilibration kinetics originating from the dynamical asymmetry between fast-diffusing HP1

### PCH condensate establishment kinetics govern focus morphology

In a wild-type context, an additional factor relevant to the coupling between chromatin and HP1-based phase separation *in nucleo* is the highly dynamic nature of the methylated chromatin landscape. This may arise from the *de novo* establishment of H3K9me2/3 domains in embryogenesis and differentiation, and the subsequent maintenance or removal of these marks during the cell cycle (14). To address the effects of H3K9me2/3 establishment on the assembly kinetics of HP1 condensates, we investigated the formation of PCH compartments during early fly embryogenesis. In Drosophila, most of the heterochromatin is localized at the pericentromeric regions, forming large, ∼20-45 Mbp-wide, contiguous H3K9me2/3 domains on each chromosome, encompassing almost 30% of the genome (52). During embryogenesis, the level of methylated chromatin increases from fertilization through nuclear cycle 14 (NC14), but visible HP1a foci first appear in cycle 11, likely reflecting the threshold of H3K9 methylation needed to nucleate HP1a condensates. In NC11-14, HP1a-PCH condensates become increasingly-large, concomitantly with the increasing length of each cell cycle (24).

We thus introduced a minimal description of H3K9me2/3 establishment in which, starting from a fully unmethylated initial state, each pericentromeric locus may stochastically acquire the H3K9me2/3 mark with a fixed methylation rate 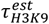. Focusing on the case of *D. melanogaster* chromosome 2 (48.8 *Mbp*, Fig.5c), we parameterized intra-nuclear HP1a and H3K9me2/3 concentrations based on mass spectrometry analysis of *D. melanogaster* embryos (53), and set 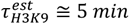 to mimic the typical evolution of methylated chromatin levels reported over the course of Drosophila NC14 (51, 54, 55) (Fig.5c, see Materials & Methods). Furthermore, in order to assess the biological relevance of our findings, we simultaneously examined the spatio-temporal dynamics of GFP-HP1a puncta across NC14 in live Drosophila embryos by means of 4D confocal microscopy (Fig.5a, see Materials & Methods). We jointly monitored the time evolution of condensate sizes and morphologies to directly juxtapose model predictions against experimental measurements over the first *τ*_*max*=_ ≅ 40 *min* of NC14.

**Figure 5:**
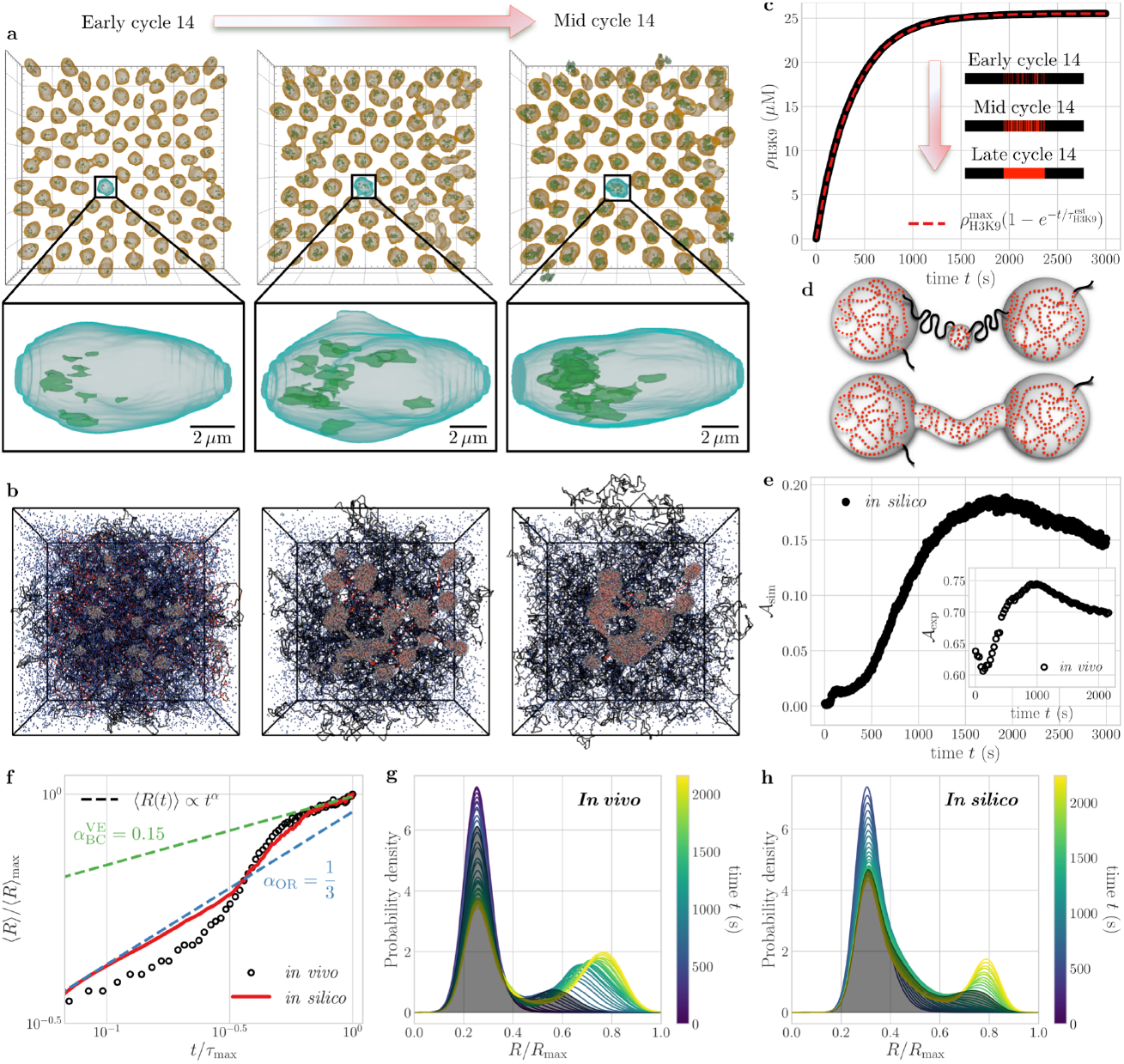
Role of H3K9me2/3 establishment kinetics: The case of Drosophila embryogenesis. (a) Surface reconstructions of HP1a foci within Drosophila nuclei during nuclear cycle (NC) 14. Time points shown are 5, 10, & 15 minutes from NC13 mitotic exit. (b) Typical kinetic pathway of the simulations over the first ∼1000 *s* of NC14 (see Movie S4). Simulation parameters as in Fig.4a but in the presence of HP1a (*J*_*HP*1_^−^_*HP*1_= 1.8 kJ/mol). (c) Modeled H3K9me2/3 establishment dynamics, with uniform methylation rate 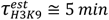 chosen to approximate the reported evolution of NC14 H3K9me2/3 levels (51). (d) Illustration of the non-local focus bridging mechanism induced by late-stage methylation events. (e) Time evolution of the focus anisotropy in both simulation (*A*_*sim*_) and *in vivo* microscopy data (*A*_*exp*_) (see SI Methods). (f) Same as (e) for the mean droplet radius ⟨*R*⟩. (g,h) Time evolution of the associated distributions of droplet size *in vivo* (g) and *in silico* (h).

Our simulations (Fig.5b) predict that the growth exponent *α* of the foci is found to display a crossover from *α* ≅ 1/3 at short times to a coarsening regime consistent with *α* ≅ 0.15 at later stages (Fig.5f), qualitatively similar to that obtained in the case of multiple competing, human-chromosome-19-like methylated domains (Fig.4c). However, this slower, competitive mode of phase separation contrasts with the faster, cooperative kinetics (α≅1/2) expected for the steady-state distribution of H3K9me2/3, which consists in one contiguous block of methylated chromatin (Fig.5c, late NC14) – much like the telocentric mouse-like chromosome model (Fig.4e). We attribute this anomalously-slow equilibration process to the dynamic propagation of H3K9me2/3, which may lead to transient stochastic patterns featuring multiple, distinct H3K9me2/3-enriched regions as methylation progresses (Fig.5c, early/mid NC14). These distinct sites in turn act as competing seeds for the nucleation of HP1 condensates. The resulting shift in the focus growth exponent from *α* ≅ 1/3 to *α* ≅ 0.15 is associated with a transition regime around *t* ∼ 10^−0.5).4^*τ*_*max*=_ ≅ 15 *min* (Fig.5f). This phenomenology does not depend strongly on the speed of H3K9me2/3 establishment (Fig.S5a,b). Moreover, it is corroborated by data analysis of live-embryo imaging, showing that the size of *in vivo* HP1 condensates exhibits very similar dynamics to those predicted by the model (Figs.5f).

Remarkably, the maturation of these HP1-PCH foci is found to be associated with an increasing loss of sphericity during the establishment of the methylated pericentromeric domain, which is evidenced in both model and experiment by a sharp rise in the mean focus anisotropy *A* during the first 20-30 min of NC14 (Fig.5e, see SI Methods) (24). The appearance of such aspherical condensates is robust to changes in the kinetics of H3K9me2/3 establishment, with slower propagation dynamics leading to a higher, more persistent anisotropy (Fig.S5c). However, it strongly contrasts with our model predictions for the coupled HP1-chromatin system in the absence of H3K9me2/3 establishment dynamics, for which *A* decays towards 0 after a 10-30 sec-long initial plateau, indicating rapid relaxation towards increasingly-round shapes in all considered scenarios (Fig.4g). We thus attribute the increase in anisotropy observed in our embryo simulations to the dynamic propagation of chromatin methylation. Indeed, as discussed further below, the creation of new H3K9me2/3-enriched regions is found to lead to the continuous nucleation of small HP1 puncta in our model, which eventually percolate into connected structures linking larger foci — associated with early-methylated chromatin domains — in a network-like fashion (Figs.5b,d).

Such complex, non-convex morphologies are consistent with *in vivo* observations at mid-NC14 (Fig.5a-e). Thus, we speculate that such structures may similarly develop *in nucleo* as the system proceeds towards full PCH establishment and maturation. This hypothesis may be supported by the analysis of the time evolution of the focus size distributions, which reveals the presence of a significant population of small HP1-PCH condensates that persists throughout most of NC14 both *in vivo* and *in silico* (Figs.5g-h). This persistent population of small puncta contrasts with the rapid disappearance of the smaller foci obtained *in silico* in the case of both pure HP1 (Fig.4f) and static methylated chromatin domains (Figs.4b,d). Furthermore, it is found to be slowly superseded by a distinct population of significantly-larger foci as the cell cycle progresses (Figs.5g-h). Therefore, this persistent HP1 population *in nucleo* could be compatible with the continuous formation of HP1-PCH condensates around newly-formed H3K9me2/3 regions, which putatively recedes as one approaches the establishment of full methylation and H3K9me2/3 nucleation subsides. At these later stages, which closely match the regime of viscoelastic-BC-like focus growth (Fig.5f), the *in vivo* condensates are found to slowly relax towards sphericity — also consistent with our model predictions (Fig.5e).

Together, these observations suggest that the dynamic establishment of H3K9me2/3 marks further promotes anomalously-slow HP1 phase separation kinetics by introducing a competition between nucleated HP1 foci within individual pericentromeres, and may lead to the development of long-lived aspherical, network-like morphologies compatible with those observed in live Drosophila embryos.

## DISCUSSION AND CONCLUSION

In this study we have used generic biophysical modeling to investigate how the propensity of architectural chromatin-binding proteins such as HP1 to undergo phase separation may be impacted by heterotypic, specific interactions with the chromatin fiber, and may in turn affect both chromosome structure and mobility. In particular, we demonstrate that the equilibrium and kinetic features displayed by *in nucleo* condensates of chromatin-binding proteins may quantitatively and qualitatively deviate from the classical hallmarks of single-component phase separation in several key aspects (56). First, our model predicts that, *in vivo*, HP1 puncta may stably form around H3K9me2/3-enriched genomic regions at physiologically-relevant concentrations of HP1, which are significantly lower than required in *in vitro* assays (Fig.3b). Such “polymer-assisted condensation” (57) emerges from the *indirect* stabilization of HP1-HP1 contacts caused by the *direct* affinity interactions between HP1 and H3K9me2/3 nucleosomes. This heterogeneous nucleation process is fully consistent with other theoretical models of self-attracting, diffusing particles – like RNA Pol II or the pioneer transcription factor Kfl4 – which may also display a specific affinity for the chromatin polymer (30, 58–60). More generally, this phenomenon is a specific illustration of polyphasic linkage (61, 62) where ligands may regulate the phase separation of scaffold molecules. In our case, methylated chromatin can be viewed as a ligand that facilitates the condensation of scaffold HP1s, as it preferentially binds to the dense scaffold (HP1) phase to maximize heterotypic interactions while maintaining high levels of homotypic interactions (63).

Second, we show that, *in vivo*, the nucleoplasmic density of HP1 in the dilute background phase is generally not constant, but instead depends on the total HP1 level (Fig.3c). When HP1 molecules are present at sub-stoichiometric levels compared to methylated chromatin sites, nucleoplasmic HP1 is buffered at a fixed value significantly lower than that measured *in vitro*. This depletion of HP1 in the dilute phase is more pronounced for stronger HP1-H3K9me2/3 interactions. Conversely, when HP1 dimers are in excess of H3K9me2/3, we predict a sub-linear increase of nucleoplasmic levels with the global HP1 concentration, associated with an anomalously-slow growth of the PCH compartment (Fig.3d). A similar increase of the nucleoplasmic concentration was recently observed in mouse embryonic fibroblast when overexpressing HP1α (28), but also in HeLa cells for several proteins like NPM1 involved in the formation of other LLPS-based condensates such as nucleoli (29). Although such effects are frequently attributed to compartmentalization driven by non-cooperative “bridger” proteins (see below) (1), our findings reveal that this behavior can also be displayed by self-interacting protein-based condensates in the presence of heterotypic interactions – in agreement with the generic properties of multicomponent phase separation (29, 64).

Third, our analysis demonstrates that, *in vivo*, the kinetics of condensate coarsening may significantly deviate from the standard OR and BC (Fig.4) and strongly depend on the linear organization of methylated chromatin along the genome, thus enabling dynamic control of the 3D PCH organization via its 1D (genomic) patterning along the polymer (65, 66). Condensate formation around long contiguous H3K9me2/3 regions is anomalously-fast (Fig.4e), and may allow the rapid recompaction of large methylated domains such as pericentromeres in late mitosis/early interphase, after the disassembly of HP1-PCH foci during mitotic prophase (24). On the contrary, the coarsening of scattered (Fig.4c) or establishing (Fig.5f) H3K9me2/3 domains is anomalously-slow and dominated by viscoelastic Brownian coalescence. These situations are consistent with the general concept of viscoelastic phase separation (102,103), in which strong asymmetries in the mobilities of the mixture components may lead to the formation of long-lived micro- or network-like phases. Interestingly, such inhibited equilibration kinetics corroborate *in vivo* measurements for HP1 (43) and FUS (46) proteins as well as *in silico* studies of nucleolus formation (48), and appear to generically characterize liquid droplets embedded in a polymer network – regardless of whether the heterotypic interactions with chromatin are attractive (48) or repulsive (46, 67). These slow coalescence dynamics may further explain why, in cycling cells, all methylated chromatin regions – initially spatially-dispersed after mitosis – generally do not colocalize into one single macro-phase, but rather form several meta-stable micro-compartments, whose fusion may be too slow to be achieved within one cell cycle (34). Our prediction is that post-replicative cells may exhibit fewer and larger PCH condensates, as has already been reported in oncogene-induced senescent fibroblasts (34) and in mouse rod photoreceptors (35).

Together, these conclusions evidence the limitations of interpreting potential intra-cellular observations of endogenous phase separation based on simple comparisons to the thermodynamic and kinetic properties of single-component phases. In this context, it is enlightening to compare our results to those of previous “bridger”-based descriptions of compartment formation such as the string-and-binders-switch model (32, 34, 65). Such approaches, which neglect the effects of HP1-HP1 homotypic interactions (Fig.S2a), are found to display many quantitative and qualitative differences with our model (Figs.S2-S3). For instance, in the “bridging-like” model, the variations in nucleoplasmic HP1 density exhibit the same qualitative trends as in our case (Fig.S2d), but the increase with HP1 concentration in the regime of stoichiometric excess is significantly more pronounced (Fig.3c). Another notable distinction between the two models lies in the interplay between HP1 and PCH condensation.

In the “bridging-like” model, both processes are tightly coupled and occur near-simultaneously (Figs.S2c,S3a), with a strongly-cooperative theta-collapse of the long H3K9me2/3 domains leading to high polymer compaction (Fig.S3a), as well as a dramatic reduction in chromatin mobility (Fig.S3c). Conversely, we observe that the nucleation of HP1 condensates at H3K9me2/3 regions may occur at much lower densities than those at which maximal PCH compaction is achieved (Figs.3b,f), which gives rise to a more diffuse transition between the dilute coiled and dense globular states. Furthermore, the chromatin configurations predicted by our model are generally less compact (∼two-fold, Figs.S3a,d) and more dynamic (∼five-fold, Figs.S3c,f) than in the “bridging-like” model which may also exhibit coarsening dynamics with viscoelastic brownian coalescence but at much slower speed (Fig.S3g,h).

Therefore, our results demonstrate that the composition dependence of nucleoplasmic HP1 levels and the presence of a switch-like coil-to-globule transition, which have been previously presented as irreconcilable with LLPS behavior (28), are actually compatible with both “bridging-like”- and LLPS-based mechanisms of chromatin condensation. Furthermore, it is worth pointing out that these two processes are not necessarily mutually-exclusive. Indeed, the biophysical properties of condensates are known to evolve over the course of the cell cycle and cellular differentiation, displaying liquid-like properties during initial establishment that transition into more static, gel-like material states consistent with maintenance (24, 68). This crossover could reflect the differential regulation of HP1 homotypic and heterotypic affinities by various post-translational modifications (69), as well as potential chromatin cross-linking by HP1 binding partners (70), which could conceivably enable endogeneous HP1 to exhibit a full spectrum of *in vivo* behaviors in-between those of the “bridger” and cooperative binding models. In biological situations like early embryogenesis, where PCH needs to be mobile and (re)organized on a large scale during the initial establishment of the condensate, dynamic LLPS-like properties may be required and functionally desirable. However, in more differentiated cells, a constrained, “bridging”-like mode of PCH segregation may be more advantageous to ensure the higher compaction and complete silencing of methylated genomic regions, or to provide specific mechanical or optical properties to the nucleus (71, 72). Indeed, HP1 mobility inside PCH has been shown to be tightly regulated, and to gradually decrease as embryogenesis progresses (24).

In addition to the magnitude of HP1-associated affinities, our analysis of early fly embryogenesis shows that the spatio-temporal dynamics of HP1 condensates also depends on the time evolution of the H3K9me2/3 mark distribution (Fig.5). In our work, to focus on HP1 condensate coarsening, we considered simplified, stochastic kinetics of H3K9me2/3 domain formation (Fig.5c, Fig.S5). More realistically, the establishment of H3K9me2/3 domains may rely on a feed-forward spreading mechanism, whereby lysine methyltransferases (KMTs) like Su(var)3-9, SetDb1 or G9a propagate methylation marks from recruitment sites via auto-catalytic reader-writer processes (14, 73, 74). As the nuclear concentration of KMTs is usually low (53), the rapid establishment of PCH domains during development would require an efficient long-range spreading activity for KMTs (75). Therefore, the spatial organization of chromatin around KMT nucleation sites may play an important role in the regulation of such activity (76). The recruitment of HP1 by nascent H3K9me2/3 regions, and the subsequent 3D compaction of methylated chromatin, could facilitate the long-range spreading of methylation marks through a positive feedback loop based on the interplay between HP1 recruitment and chromatin compaction, which may accelerate the establishment of epigenetic patterns (77–79). While this amplification loop may be essential to facilitate the spreading of PCH, the very good agreement between our predictions and experiments (Figs.5e-h) indicates that the large-scale coarsening dynamics of HP1 condensates are not strongly limited by the underlying kinetics of H3K9me2/3 establishment. Instead, our results would suggest the aspherical morphology of such condensates as a generic experimental signature of the dynamic character of the epigenetic landscape being established (Figs.5d-e).

To conclude, we have developed a generic quantitative framework to investigate the phase separation of architectural chromatin-binding proteins *in nucleo*, and contextualized our investigations to the formation of PCH condensates in higher eukaryotes. However, to perform such a systematic analysis, we had to resort to several simplifying assumptions. First, our lattice-gas model emulates, in the absence of the polymer, a coarsening dynamics dominated by OR over BC. However, the very few quantitative studies investigating the *in vitro* coarsening of proteins involved in the formation of *in vivo* biomolecular condensates suggests that BC might be the dominant mode (8). This may impact the early coarsening regime of scattered H3K9me2/3 domains (Fig.4c) where OR still dominates, before viscoelastic BC drives the kinetics. Second, we limited our analysis to a finite-size system consisting of a single chromosome. Our results in the case of the human (Fig.4) and Drosophila (Fig.5) nuclei suggest that the presence of other chromosomes – and thus, of other competing H3K9me2/3 regions – could add an additional layer of viscoelastic coarsening dynamics driven by inter-chromosomal diffusion kinetics. Such very slow processes (80) could prohibit the fusion of inter-chromosomal condensates, except in cases where the majority of H3K9me2/3 regions are co-localized post-mitosis, as in the Rabl configuration observed during fly embryogenesis (81). Third, we did not explicitly consider the dimerization thermodynamics of HP1, the finite interaction valency of HP1 with chromatin (82, 83), or the non-specific interactions between HP1 and DNA (19, 23, 84, 85). These additional components and effects could possibly alter the impact of HP1 on PCH compaction (60, 86) *in vivo* - especially for endogenous HP1 levels close to the dimer dissociation constant - but the overall phenomenology is likely to be conserved (Fig. S4). Finally, we restricted our work to a binary HP1-H3K9me2/3 system, neglecting other interaction partners such as RNA or HP1 binding factors that may interfere with or facilitate the formation of PCH condensates, which could serve to further enhance the complexity of the corresponding multicomponent phase diagram (87, 88).

Future studies will thus have to integrate such ingredients, coupled to a more precise description of H3K9me2/3 establishment, in order to fully characterize the spatio-temporal dynamics of PCH condensates in various biological contexts – such as the DNA damage repair of methylated chromatin loci (89) – as well as to investigate the important role of HP1 in structuring non-pericentromeric regions (90, 91). Nonetheless, the examination of the transferability of our minimal model to other architectural proteins constitutes a promising avenue of research, in light of its remarkable ability to correctly capture the *in vivo* features of HP1-based PCH condensate establishment in embryogenesis. In this framework, a natural potential candidate is PRC1, which plays a crucial role in the developmental regulation of Polycomb target genes (92) and has similarly been shown to exhibit LLPS-like properties *in vitro* (93, 94).

## ACKNOWLEDGEMENTS

We are grateful to members of the Karpen and Jost labs for fruitful discussions and Geoff Fudenberg and Aurèle Piazza for a critical reading of the manuscript. Part of this work was initiated during the KITP Program: Biological Physics of Chromosomes 2020 (supported by the National Science Foundation under Grant No. NSF PHY-1748958 and NIH Grant No. R25GM067110). Work in the Jost lab is funded by Agence Nationale de la Recherche (ANR-18-CE12-0006-03; ANR-18-CE45-0022-01). The Karpen lab acknowledges National Institute of Health (R35GM139653) and Volkswagen Stiftung (98196). We thank PSMN (Pôle Scientifique de Modélisation Numérique) and CBP (Centre Blaise Pascal) of the ENS de Lyon for computing resources.

## MATERIALS & METHODS

### Lattice-gas model of HP1-based phase separation

Individual HP1 dimers were represented as spherical beads with effective diameter *δ* residing on the vertices of a face-centered cubic lattice *L*. Multivalent affinity interactions between proximal dimers were described by a nearest-neighbor pair potential of depth *J*_*HP*1− *HP*1_,

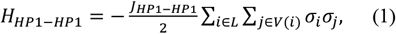

where the double sum runs over each lattice site *i* and its 12 connected neighbors *j* ∈ *V*(*i*). In Eq. (1), the occupancy number *σ*_*i*_ equals 1 if a dimer is present at site *i* and 0 otherwise, reflecting the fact that each lattice site may contain at most a single HP1 dimer. The lattice is comprised of a finite number *N* of vertices, and periodic boundary conditions were used to mimic potential cross-interactions with vicinal intra-nuclear regions.

Simulations were initialized by uniformly distributing a number *N*_*HP*1_ of HP1 dimers on the lattice in a random arrangement, and were evolved through standard kinetic Monte Carlo (MC) rules at fixed temperature *T* = 300 *K* (95). In this context, the critical point *J*_*crit*_ of the lattice-gas model for FCC lattice is *J*_*crit*_≃ 0.4 *k*_6_*T* ≃ 1 kJ/mol (96), with *k*_*B*_ the Boltzmann constant, which marks the dimer-dimer interaction threshold below which HP1 may not spontaneously phase separate at any concentration. Each MC step then consists of an arbitrary number *N*_*trial*_ of trial moves, in which a constituent HP1 dimer is first selected at random, whose position on the lattice we denote by *p*. The particle is then displaced to a random neighboring site *q* ∈ *V*(*p*), with an acceptance probability provided by the Metropolis criterion associated with the Hamiltonian in Eq. (1). In the event that site *q* is also occupied by another HP1 dimer, we implement a particle exchange protocol between sites *p* and *q* with an acceptance probability of 1, reflecting the invariance of Eq. (1) to such molecular swap moves. These evolution rules ensure that the system emulates Cahn-Hilliard-Cook (also known as *Model B*) dynamics in the continuum limit (97), and is therefore suitable to describe phase separation kinetics in the case of negligible hydrodynamic interactions (8) – an assumption consistent with the hydrodynamic screening approximation commonly used in numerical simulations of chromatin-based processes *in nucleo* (98).

In this framework, the overall molecular level *ρ*_*HP*1_ of HP1 is governed by the fraction *N*_*HP*1_/*N* of occupied lattice sites, which remains fixed over the course of the simulations, and is such that (*N*_*HP*1_/*N*) ≲ 0.1 for all systems considered here. Since we here mostly focus on the density regime *ρ*_*HP*1_ ≳ *K*_*d*_, with *K*_*d*_ ≅ 1 *μM* the HP1 dimer dissociation constant, we assume that the majority of HP1 is present in dimerized form, and neglect the population of free HP1 monomers as a first approximation. In this case, the molar concentration of HP1 simply reads as *ρ*_*HP*1_ = 2(*N*_*HP*1_/*N*)/(*N*_*A*_*v*_*site*_), with *N*_*A*_ the Avogadro constant and 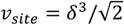 the effective volume of each lattice site. We set *δ* = 20 *nm*, consistent with the approximate radius of gyration 2*R*_*gyr*_≅ 15 *nm* of the extended conformation of HP1α dimers (23), which has been shown to be generally prevalent in phase-separated HP1 assemblies (21, 23). *R*_*gyr*_ was estimated from the corresponding SAXS distance distribution *P*(*r*) (23) via (99)

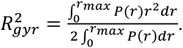

### Polymer model of chromatin

Chromatin was described as a self-avoiding, semi-flexible polymer chain comprised of *N*_*chr*_ monomers, represented as spherical beads of diameter *δ* residing on the same lattice *L* as the HP1 dimers. The bending rigidity of the chromatin fiber was incorporated via a standard angular potential of stiffness *κ*,

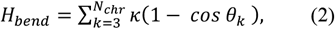

where *θ*_*K*_ denotes the angle formed by the triplet of adjacent monomers (*k* − 2, *k* − 1, *k*). Assuming a standard chromatin compaction value of 50 bp/nm, the effective chain diameter *δ* = 20 *nm* commensurate with the typical HP1 dimer size implies that each monomer within the chain encapsulates a genomic locus of approximate length *ϑ* = 1 *kbp*. Accordingly, we set the polymer bending modulus to *κ* = 3.217 *k*_*B*_*T*, corresponding to a Kuhn length *l*_*K*_ = 100 *nm* consistent with the predictions of coarse-grained chromatin models at similar levels of spatial resolution (100, 101).

Each monomer was considered to be in one of two chromatin states, respectively depicted in red and black in Figs.3-5, in order to differentiate genomic domains bearing the H3K9me2/3 histone modifications (red) from euchromatic regions (black). The chromodomain-mediated binding affinity of HP1 for H3K9-methylated histone tails was accounted for through a short-ranged attractive potential of depth *J*_*HP*1−*H*3*K*9_,

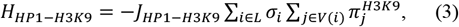

where the occupancy number 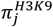 quantifies the number of H3K9-methylated loci present at a given lattice site *j*. In the model, one H3K9 monomer can interact with, at most, 13 HP1 which is consistent with the maximal number of H3K9me2/3 residues (∼10) present in a fully methylated 1-kbp long region.

Whole chromosome simulations were performed starting from dense, random and unknotted initial configurations, and were similarly evolved via a kinetic MC scheme, as detailed in previous work (95). In this case, each MC step consists of a number *N*_*chr*_ of local trial displacements involving individual monomers, including both translation and reptation moves (38). Polymer acceptance rates were computed from Eqs. (2) and (3) based on the Metropolis rule (95), and Eq. (3) was also incorporated into the Hamiltonian in Eq. (1) for the determination of HP1 acceptance probabilities in coupled chromatin-HP1 simulations. Simulation and analysis codes are available at https://github.com/physical-biology-of-chromatin/LatticePoly.

### Live embryo imaging & analysis

Flies homozygous for the expression of ectopic GFP-HP1a on the second chromosome (+, GFP-HP1a/CyO; +)(24) were allowed to lay embryos on apple juice plates supplemented with yeast paste at 25°C. Embryos were collected by hand and dechorionated in 50% bleach before being mounted for live imaging. Embryos were imaged using a Zeiss 880 Airyscan microscope with a 63x/1.4 oil immersion objective. Time-lapse image stacks were collected every 30 seconds with a Z-spacing of 0.36μm. Imaging at least part of nuclear cycle 13 (NC 13) allowed the start of NC14 interphase (time=0 sec) to be clearly defined as the first time point where circular nuclei appear following mitosis. All movies were Airyscan-processed using Zen 2.3 software and the first 72 time points of NC 14 (*τ*_*max*_ ≃36 minutes) were analyzed using Arivis software. Briefly, HP1a foci were segmented from the overall nuclear signal using an intensity threshold > 75% of the fluorescence intensity and a lower size threshold of 0.03 μm^3^ corresponding to approximately 10 voxels (0.085 μm x 0.085 μm x 0.36 μm). All HP1a segments found to be connected in 3-dimensional space were merged into a single segment, and population statistics were calculated for each time point. Imaging data are available at doi:10.6078/D1TQ67.

## REFERENCES

1. P. Bhat, D. Honson, M. Guttman, Nuclear compartmentalization as a mechanism of quantitative control of gene expression. Nat. Rev. Mol. Cell Biol. 22, 653–670 (2021).

2. M. Lisby, R. Rothstein, U. H. Mortensen, Rad52 forms DNA repair and recombination centers during S phase. Proc. Natl. Acad. Sci. U.S.A. 98, 8276–8282 (2001).

3. A. Papantonis, P. R. Cook, Transcription factories: genome organization and gene regulation. Chem. Rev. 113, 8683–8705 (2013).

4. B. R. Sabari, et al., Coactivator condensation at super-enhancers links phase separation and gene control. Science 361 (2018).

5. N. Saner, et al., Stochastic association of neighboring replicons creates replication factories in budding yeast. J. Cell Biol. 202, 1001–1012 (2013).

6. E. R. Gibney, C. M. Nolan, Epigenetics and gene expression. Heredity 105, 4–13 (2010).

7. Y. Shin, C. P. Brangwynne, Liquid phase condensation in cell physiology and disease. Science 357 (2017).

8. J. Berry, C. P. Brangwynne, M. Haataja, Physical principles of intracellular organization via active and passive phase transitions. Rep. Prog. Phys. 81, 046601 (2018).

9. A. A. Hyman, C. A. Weber, F. Jülicher, Liquid-liquid phase separation in biology. Annu. Rev. Cell Dev. Biol. 30, 39–58 (2014).

10. C. P. Brangwynne, P. Tompa, R. V. Pappu, Polymer physics of intracellular phase transitions. Nature Physics 11, 899–904 (2015).

11. D. T. McSwiggen, M. Mir, X. Darzacq, R. Tjian, Evaluating phase separation in live cells: diagnosis, caveats, and functional consequences. Genes Dev. 33, 1619–1634 (2019).

12. F. Erdel, K. Rippe, Formation of Chromatin Subcompartments by Phase Separation. Biophysical Journal 114, 2262–2270 (2018).

13. M. L. Heltberg, J. Miné-Hattab, A. Taddei, A. M. Walczak, T. Mora, Physical observables to determine the nature of membrane-less cellular sub-compartments. Elife 10 (2021).

14. R. C. Allshire, H. D. Madhani, Ten principles of heterochromatin formation and function. Nat. Rev. Mol. Cell Biol. 19, 229–244 (2018).

15. J. C. Eissenberg, S. C. Elgin, The HP1 protein family: getting a grip on chromatin. Curr. Opin. Genet. Dev. 10, 204–210 (2000).

16. D. L. Mendez, R. E. Mandt, S. C. R. Elgin, Heterochromatin Protein 1a (HP1a) partner specificity is determined by critical amino acids in the chromo shadow domain and C-terminal extension. J. Biol. Chem. 288, 22315–22323 (2013).

17. A. J. Bannister, et al., Selective recognition of methylated lysine 9 on histone H3 by the HP1 chromo domain. Nature 410, 120–124 (2001).

18. A. Thiru, et al., Structural basis of HP1/PXVXL motif peptide interactions and HP1 localisation to heterochromatin. EMBO J. 23, 489–499 (2004).

19. D. Canzio, et al., Chromodomain-mediated oligomerization of HP1 suggests a nucleosome-bridging mechanism for heterochromatin assembly. Mol. Cell 41, 67–81 (2011).

20. S. Machida, et al., Structural Basis of Heterochromatin Formation by Human HP1. Mol. Cell 69, 385–397.e8 (2018).

21. A. Kumar, H. Kono, Heterochromatin protein 1 (HP1): interactions with itself and chromatin components. Biophysical Reviews 12, 387–400 (2020).

22. A. P. Latham, B. Zhang, Consistent Force Field Captures Homologue-Resolved HP1 Phase Separation. J. Chem. Theory Comput. 17, 3134–3144 (2021).

23. A. G. Larson, et al., Liquid droplet formation by HP1α suggests a role for phase separation in heterochromatin. Nature 547, 236–240 (2017).

24. A. R. Strom, et al., Phase separation drives heterochromatin domain formation. Nature 547, 241–245 (2017).

25. M. M. Keenen, et al., HP1 proteins compact DNA into mechanically and positionally stable phase separated domains. Elife 10 (2021).

26. T. Cheutin, et al., Maintenance of stable heterochromatin domains by dynamic HP1 binding. Science 299, 721–725 (2003).

27. L. Wang, et al., Histone Modifications Regulate Chromatin Compartmentalization by Contributing to a Phase Separation Mechanism. Mol. Cell 76, 646–659.e6 (2019).

28. F. Erdel, et al., Mouse Heterochromatin Adopts Digital Compaction States without Showing Hallmarks of HP1-Driven Liquid-Liquid Phase Separation. Mol. Cell 78, 236–249.e7 (2020).

29. J. A. Riback, et al., Composition-dependent thermodynamics of intracellular phase separation. Nature 581, 209–214 (2020).

30. M. Ancona, C. A. Brackley, Simulating the chromatin-mediated phase separation of model proteins with multiple domains. Biophys. J. 121, 2600–2612 (2022).

31. M. Barbieri, et al., Complexity of chromatin folding is captured by the strings and binders switch model. Proc. Natl. Acad. Sci. U. S. A. 109, 16173–16178 (2012).

32. Q. MacPherson, B. Beltran, A. J. Spakowitz, Bottom–up modeling of chromatin segregation due to epigenetic modifications. Proc. Natl. Acad. Sci. U.S.A. 115, 12739–12744 (2018).

33. A. Esposito, et al., Polymer physics reveals a combinatorial code linking 3D chromatin architecture to 1D chromatin states. Cell Rep. 38, 110601 (2022).

34. S. Sati, et al., 4D Genome Rewiring during Oncogene-Induced and Replicative Senescence. Mol. Cell 78, 522–538.e9 (2020).

35. M. Falk, et al., Heterochromatin drives compartmentalization of inverted and conventional nuclei. Nature 570, 395–399 (2019).

36. A. Z. Abdulla, H. Salari, M. M. C. Tortora, C. Vaillant, D. Jost, 4D epigenomics: deciphering the coupling between genome folding and epigenomic regulation with biophysical modeling. Curr. Opin. Genet. Dev. 79, 102033 (2023).

37. M. R. Williams, Y. Xiaokang, N. A. Hathaway, D. Kireev, A simulation model of heterochromatin formation at submolecular detail. iScience 25, 104590 (2022).

38. S. K. Ghosh, D. Jost, How epigenome drives chromatin folding and dynamics, insights from efficient coarse-grained models of chromosomes. PLoS Comput. Biol. 14, e1006159 (2018).

39. D. H. Rothman, S. Zaleski, Lattice-gas models of phase separation: interfaces, phase transitions, and multiphase flow. Reviews of Modern Physics 66, 1417–1479 (1994).

40. S. F. Banani, H. O. Lee, A. A. Hyman, M. K. Rosen, Biomolecular condensates: organizers of cellular biochemistry. Nat. Rev. Mol. Cell Biol. 18, 285–298 (2017).

41. J.-M. Choi, A. S. Holehouse, R. V. Pappu, Physical Principles Underlying the Complex Biology of Intracellular Phase Transitions. Annu. Rev. Biophys. 49, 107–133 (2020).

42. M. M. Tortora, H. Salari, D. Jost, Chromosome dynamics during interphase: a biophysical perspective. Curr. Opin. Genet. Dev. 61, 37–43 (2020).

43. Y. Gao, M. Han, S. Shang, H. Wang, L. S. Qi, Interrogation of the dynamic properties of higher-order heterochromatin using CRISPR-dCas9. Mol. Cell 81, 4287–4299.e5 (2021).

44. I. Solovei, K. Thanisch, Y. Feodorova, How to rule the nucleus: divide et impera. Curr. Opin. Cell Biol. 40, 47–59 (2016).

45. D. Jost, P. Carrivain, G. Cavalli, C. Vaillant, Modeling epigenome folding: formation and dynamics of topologically associated chromatin domains. Nucleic Acids Res. 42, 9553–9561 (2014).

46. D. S. W. Lee, N. S. Wingreen, C. P. Brangwynne, Chromatin Mechanics Dictates Subdiffusion and Coarsening Dynamics of Embedded Condensates. Nature Physics 17, 531–538 (2021).

47. Y. Shin, et al., Liquid Nuclear Condensates Mechanically Sense and Restructure the Genome. Cell 176, 1518 (2019).

48. Y. Qi, B. Zhang, Chromatin network retards nucleoli coalescence. Nat. Commun. 12, 6824 (2021).

49. A. J. Webster, M. E. Cates, Stabilization of Emulsions by Trapped Species. Langmuir 14, 2068 (1998).

50. W. Xu, Z. Lan, B. Peng, R. Wen, X. Ma, Evolution of transient cluster/droplet size distribution in a heterogeneous nucleation process. RSC Adv. 4, 31692–31699 (2014).

51. K. H.-C. Wei, C. Chan, D. Bachtrog, Establishment of H3K9me3-dependent heterochromatin during embryogenesis in Drosophila miranda. eLife 10 (2021).

52. C. D. Smith, S. Shu, C. J. Mungall, G. H. Karpen, The Release 5.1 annotation of Drosophila melanogaster heterochromatin. Science 316, 1586–1591 (2007).

53. J. Bonnet, et al., Quantification of Proteins and Histone Marks in Drosophila Embryos Reveals Stoichiometric Relationships Impacting Chromatin Regulation. Dev. Cell 51, 632–644.e6 (2019).

54. K. Yuan, P. H. O’Farrell, TALE-light imaging reveals maternally guided, H3K9me2/3-independent emergence of functional heterochromatin in Drosophila embryos. Genes Dev. 30, 579–593 (2016).

55. C. A. Seller, C.-Y. Cho, P. H. O’Farrell, Rapid embryonic cell cycles defer the establishment of heterochromatin by Eggless/SetDB1 in. Genes Dev. 33, 403–417 (2019).

56. D. Bracha, M. T. Walls, C. P. Brangwynne, Probing and engineering liquid-phase organelles. Nat. Biotechnol. 37, 1435–1445 (2019).

57. J.-U. Sommer, H. Merlitz, H. Schiessel, Polymer-Assisted Condensation: A Mechanism for Hetero-Chromatin Formation and Epigenetic Memory. Macromolecules 55, 11, 4841–4851 (2022).

58. A. Pancholi, et al., RNA polymerase II clusters form in line with surface condensation on regulatory chromatin. Mol. Syst. Biol. 17, e10272 (2021).

59. J. A. Morin, et al., Surface condensation of a pioneer transcription factor on DNA https://doi.org/10.1101/2020.09.24.311712.

60. I. Malhotra, B. Oyarzún, B. M. Mognetti, Unfolding of the chromatin fiber driven by overexpression of noninteracting bridging factors. Biophys. J. 120, 1247–1256 (2021).

61. K. M. Ruff, F. Dar, R. V. Pappu, Polyphasic linkage and the impact of ligand binding on the regulation of biomolecular condensates. Biophys. Rev. 2, 021302 (2021).

62. J. Wyman, S. J. Gill, Ligand-linked phase changes in a biological system: applications to sickle cell hemoglobin. Proc. Natl. Acad. Sci. U. S. A. 77, 5239–5242 (1980).

63. K. M. Ruff, F. Dar, R. V. Pappu, Ligand effects on phase separation of multivalent macromolecules. Proc. Natl. Acad. Sci. U. S. A. 118 (2021).

64. J.-M. Choi, F. Dar, R. V. Pappu, LASSI: A lattice model for simulating phase transitions of multivalent proteins. PLoS Comput. Biol. 15, e1007028 (2019).

65. E. W. Martin, et al., Valence and patterning of aromatic residues determine the phase behavior of prion-like domains. Science 367, 694–699 (2020).

66. X. Zeng, K. M. Ruff, R. V. Pappu, Competing interactions give rise to two-state behavior and switch-like transitions in charge-rich intrinsically disordered proteins. Proc. Natl. Acad. Sci. U. S. A. 119, e2200559119 (2022).

67. Y. Zhang, D. S. W. Lee, Y. Meir, C. P. Brangwynne, N. S. Wingreen, Mechanical Frustration of Phase Separation in the Cell Nucleus by Chromatin. Phys. Rev. Lett. 126, 258102 (2021).

68. I. Eshghi, J. A. Eaton, A. Zidovska, Interphase Chromatin Undergoes a Local Sol-Gel Transition upon Cell Differentiation. Phys. Rev. Lett. 126, 228101 (2021).

69. G. LeRoy, et al., Heterochromatin protein 1 is extensively decorated with histone code-like post-translational modifications. Mol. Cell. Proteomics 8, 2432–2442 (2009).

70. M. Jagannathan, R. Cummings, Y. M. Yamashita, The modular mechanism of chromocenter formation in Drosophila (2019) https://doi.org/10.7554/eLife.43938 (June 22, 2022).

71. G. Gerlitz, The Emerging Roles of Heterochromatin in Cell Migration. Front Cell Dev Biol 8, 394 (2020).

72. Y. Feodorova, M. Falk, L. A. Mirny, I. Solovei, Viewing Nuclear Architecture through the Eyes of Nocturnal Mammals. Trends Cell Biol. 30, 276–289 (2020).

73. P. Cruz-Tapias, P. Robin, J. Pontis, L. D. Maestro, S. Ait-Si-Ali, The H3K9 Methylation Writer SETDB1 and its Reader MPP8 Cooperate to Silence Satellite DNA Repeats in Mouse Embryonic Stem Cells. Genes 10 (2019).

74. A. R. Cutter DiPiazza, et al., Spreading and epigenetic inheritance of heterochromatin require a critical density of histone H3 lysine 9 tri-methylation. Proc. Natl. Acad. Sci. U. S. A. 118 (2021).

75. M. J. Obersriebnig, E. M. H. Pallesen, K. Sneppen, A. Trusina, G. Thon, Nucleation and spreading of a heterochromatic domain in fission yeast. Nat. Commun. 7, 11518 (2016).

76. D. Jost, C. Vaillant, P. Meister, Coupling 1D modifications and 3D nuclear organization: data, models and function. Curr. Opin. Cell Biol. 44, 20–27 (2017).

77. D. Jost, C. Vaillant, Epigenomics in 3D: importance of long-range spreading and specific interactions in epigenomic maintenance. Nucleic Acids Res. 46, 2252–2264 (2018).

78. D. Michieletto, E. Orlandini, D. Marenduzzo, A Polymer Model with Epigenetic Recolouring Reveals a Pathway for the de novo Establishment and 3D organisation of Chromatin Domains. Phys. Rev. X 6, 041047 (2016).

79. S. H. Sandholtz, Q. MacPherson, A. J. Spakowitz, Physical modeling of the heritability and maintenance of epigenetic modifications. Proc. Natl. Acad. Sci. U. S. A. 117, 20423–20429 (2020).

80. A. Rosa, R. Everaers, Structure and dynamics of interphase chromosomes. PLoS Comput. Biol. 4, e1000153 (2008).

81. W. F. Marshall, A. F. Dernburg, B. Harmon, D. A. Agard, J. W. Sedat, Specific interactions of chromatin with the nuclear envelope: positional determination within the nucleus in Drosophila melanogaster. Mol. Biol. Cell 7, 825–842 (1996).

82. G. Nishibuchi, J.-I. Nakayama, Biochemical and structural properties of heterochromatin protein 1: understanding its role in chromatin assembly. J. Biochem. 156, 11–20 (2014).

83. D. Canzio, A. Larson, G. J. Narlikar, Mechanisms of functional promiscuity by HP1 proteins. Trends Cell Biol. 24, 377–386 (2014).

84. S. Kilic, A. L. Bachmann, L. C. Bryan, B. Fierz, Multivalency governs HP1α association dynamics with the silent chromatin state. Nature Communications 6 (2015).

85. D. Canzio, et al., A conformational switch in HP1 releases auto-inhibition to drive heterochromatin assembly. Nature 496, 377–381 (2013).

86. S. Guha, M. K. Mitra, Multivalent binding proteins can drive collapse and reswelling of chromatin in confinement. Soft Matter 19, 153–163 (2023).

87. S. Maharana, et al., RNA buffers the phase separation behavior of prion-like RNA binding proteins. Science 360, 918–921 (2018).

88. E. M. Langdon, et al., mRNA structure determines specificity of a polyQ-driven phase separation. Science 360, 922–927 (2018).

89. A. Janssen, et al., A single double-strand break system reveals repair dynamics and mechanisms in heterochromatin and euchromatin. Genes Dev. 30, 1645–1657 (2016).

90. F. Zenk, et al., HP1 drives de novo 3D genome reorganization in early Drosophila embryos. Nature 593, 289–293 (2021).

91. Y. C. G. Lee, et al., Pericentromeric heterochromatin is hierarchically organized and spatially contacts H3K9me2 islands in euchromatin. PLoS Genet. 16, e1008673 (2020).

92. B. Schuettengruber, H.-M. Bourbon, L. Di Croce, G. Cavalli, Genome Regulation by Polycomb and Trithorax: 70 Years and Counting. Cell 171, 34–57 (2017).

93. A. J. Plys, et al., Phase separation of Polycomb-repressive complex 1 is governed by a charged disordered region of CBX2. Genes Dev. 33, 799–813 (2019).

94. E. Seif, et al., Phase separation by the polyhomeotic sterile alpha motif compartmentalizes Polycomb Group proteins and enhances their activity. Nat. Commun. 11, 5609 (2020).

95. K. Binder, Monte Carlo Methods in Statistical Physics (Springer Science & Business Media, 2012).

96. P. H. Lundow, K. Markström, A Rosengren. The Ising model for the bcc, fcc and diamond lattices; a comparison. Philosophical Magazine 89, 2009–2042 (2009).

97. M. A. Katsoulakis, D. G. Vlachos, Coarse-grained stochastic processes and kinetic Monte Carlo simulators for the diffusion of interacting particles. The Journal of Chemical Physics 119, 9412–9427 (2003).

98. H. Hajjoul, et al., High-throughput chromatin motion tracking in living yeast reveals the flexibility of the fiber throughout the genome. Genome Res. 23, 1829–1838 (2013).

99. A. G. Kikhney, D. I. Svergun, A practical guide to small angle X-ray scattering (SAXS) of flexible and intrinsically disordered proteins. FEBS Lett. 589, 2570–2577 (2015).

100. J.-M. Arbona, S. Herbert, E. Fabre, C. Zimmer, Inferring the physical properties of yeast chromatin through Bayesian analysis of whole nucleus simulations. Genome Biol. 18, 81 (2017).

101. M. Socol, et al., Rouse model with transient intramolecular contacts on a timescale of seconds recapitulates folding and fluctuation of yeast chromosomes. Nucleic Acids Res. 47, 6195–6207 (2019).

102. H. Tanaka, Viscoelastic phase separation in biological cells. Communications Physics 5, 167 (2022).

103. H. Tanaka, Viscoelastic phase separation. J. Phys.: Condens. Matter 12, R207 (2000).

