## Supplementary Information for "HP1-driven phase separation recapitulates the thermodynamics and kinetics of heterochromatin condensate formation"

**This PDF file includes:**

Extended numerical methods
Figures S1 to S7
Legends for Movies S1 to S4
SI References

**Other supplementary materials for this manuscript include the following:**

Movies S1 to S4

### EXTENDED NUMERICAL METHODS

#### Model parametrization & time mapping

The length of the simulated chromatin chain is given by $N_{chr} = \frac{N_{bp}}{\vartheta}$, with $N_{bp}$ the total genomic size of the chromosome considered, and was set to $N_{chr} = 60{,}000$ monomers in the case of mouse-like and human chromosome 19 ($N_{bp} \cong 60\ Mbp$), and $N_{chr} = 48{,}800$ for *D. melanogaster* chromosome 2 ($N_{bp} \cong 48.8\ Mbp$). The lattice size $N$ was adjusted to yield a DNA density $\rho = \frac{N_{chr}}{N} \times \frac{\vartheta}{v_{site}}$ consistent with the typical value $\rho \cong 0.01\ bp/nm^3$ observed in mammalian and fly nuclei (1). $N_{HP1}$ was accordingly varied in the approximate range $[1{,}000{:}\,20{,}000]$ dimers, in order to probe a concentration regime $\rho_{HP1} \in [1\ \mu M, 15\ \mu M]$ encompassing the characteristic endogenous HP1 levels reported in mice nuclei (2).

In the case of idealized telocentric mouse chromosome 19, 10,000 consecutive monomers at one end of the polymeric chain were assigned a H3K9 state. In the case of idealized human chromosome 19, the H3K9 methylation pattern was obtained from ChIP sequencing data in primary fibroblasts (3), to which a centromere was appended between genomic positions $24.2\ Mbp$ and $28.1\ Mbp$. This amounts to a total number $N_{H3K9}{\sim}22{,}000$ of H3K9me2/3 loci within the chromatin chain. For simulations of Drosophila embryos, a time-dependent heterochromatin distribution was used, following the numerical protocol described in the next section. Note that, in our framework, we assumed that the H3K9 status of one monomer is binary (no methylation or fully methylated) and remain constant (during the whole simulations in the mouse- and human-like cases or after its establishment in the drosophila-like case, see below), hence neglecting the intrinsic dynamics of H3K9me2/3 that may be added or removed by dedicated enzymes or diluted during replication (18). For example, for cycling cells, just after mitosis, when HP1-heterochromatin condensates form, the levels of H3K9me2/3 are about 70-80% of their maximal levels (in S phase) and then gradually increase during G1 (19). Since, in our framework, being partially methylated, would be equivalent to effectively reducing the strength of HP1-H3K9 interaction (less residues in a 1-kbp monomer able to interact with HP1), it would correspond to weak dynamical change of $J_{HP1-H3K9}$ by 25%, which, from our systematic analysis of this parameter (Fig.3 & 4 of the main text), should not change the phenomenology of the system.

The polymer was systematically relaxed over $O(10^6)$ MC steps in the absence of HP1 interactions prior to any data collection in order to reach the metastable state of a crumpled polymer (1). Mean squared displacements (MSDs) were then computed in the standard fashion,

$$MSD_k(t) = < [r_k(t) - r_k(0)]^2 >,$$

with $r_k(t)$ the position of an individual monomer $k$ at time $t$, and were further averaged over all time intervals $\Delta t$ and H3K9me2/3 loci $k$ to yield $MSD_{H3K9}(\Delta t)$. Heterochromatin diffusion constants $D_{H3K9}$ and exponents $\beta_{H3K9}$ were defined as

$$MSD_{H3K9}(\Delta t) \sim D_{H3K9} \Delta t^{\beta_{H3K9}}, \qquad \text{(S1)}$$

and were obtained by fitting Eq. (S1) numerically via Powell's conjugate direction method (4). The experimental mapping of simulation times was performed by comparing the ensemble-averaged 2D MSD of individual loci to the typical diffusion coefficient $D_0 \cong 0.01\ \mu m^2/s^{0.5}$ measured in yeast and Drosophila nuclei (1, 5), which leads to an approximate correspondence of $\sim 150\ \mu s$ per MC step. Accordingly, the number of HP1 trial moves per MC step was set to $N_{trial} = N_{HP1}$, which was similarly found to yield a simulated diffusion coefficient $D_{HP1} \cong 1\ \mu m^2/s$ consistent with the characteristic values reported by fluorescence correlation spectroscopy studies of HP1 diffusion *in nucleo* (6, 7).

The density $\rho^0_{HP1\alpha}$of HP1α dimers was estimated from the experimental turbidity curve obtained for purified HP1α assays in physiological buffer conditions (2), and was defined as the HP1 concentration measured at the onset of turbidity in the system – which constitutes a proxy for the onset of phase separation (8). This quantity was evaluated as the abscissa of intersection of the tangent to the sigmoidal turbidity curve at the half-saturation point with the $y = 0$ lower asymptote, which yields $\rho^0_{HP1\alpha} \cong 7\ \mu M$ (2). For HP1a, $\rho^0_{HP1a}$ was inferred from the density $\sim 0.5\ mg/mL$ reported at the onset of droplet formation for purified HP1a at $100\ mM$ ionic strength (9), which amounts to $\rho^0_{HP1a} \cong 20\ \mu M$. Using Figs.2b,c, the corresponding dilute-phase concentrations of purified HP1α and HP1a respectively read as $\rho^{dil}_{HP1\alpha} \cong 3\ \mu M$ and $\rho^{dil}_{HP1a} \cong 10\ \mu M$.

Note that, unless specified and discussed in the main text, most of our results are robust and do not depend on the fine details of the model as illustrated by the consistency observed among the various investigated situations (systematic analysis of $\rho_{HP1}$ and $J_{HP1-H3K9}$ for two $J_{HP1-HP1}$ values [$HP1\alpha$ and $HP1a$] and two H3K9 patterns [telocentric mouse-like and human-like]).

**H3K9 establishment simulations in Drosophila embryos**

In the case of Drosophila embryos, HP1 and H3K9me2/3 concentrations were inferred from mass spectroscopy measurements performed across cellular cycles 9-15 of *D. melanogaster* embryogenesis (10). Assuming a typical nuclear volume $V_{nucl} \cong 90\ \mu m^3$, consistent with optical microscopy measurements in fly embryos (11), the corresponding HP1 and H3K9me2/3 densities respectively read as $\rho_{HP1a} \cong 8\ \mu M$ and $\rho_{H3K9} \cong 25\ \mu M$. The associated number of H3K9-methylated loci is thus $N_{H3K9} = \rho_{H3K9} \times N N_A v_{site} \times \frac{r}{2\vartheta} \sim 17{,}000$ monomers, where $r = 184$ bp is the average nucleosome repeat length in Drosophila embryos (12) and the factor 2 accounts for the two H3 tails of each histone octamer.

Methylated loci were assumed to be co-localized within a single consecutive chromatin segment of length $N_{H3K9}$ at the middle of the polymer, in order to emulate the centromeric and pericentromeric regions of *D. melanogaster* chromosome 2 (Fig.5c). The dynamic establishment of H3K9 methylation marks was modeled by simple stochastic activation of pericentromeric loci with uniform rate $\frac{1}{\tau^{est}_{H3K9}}$, starting from a fully-unmethylated state at time 0 (Fig.5c). To mimic the approximate time evolution of H3K9 methylation levels reported in Ref. (13) for nuclear cycle 14 (NC14) of *D. miranda* embryogenesis, we set $\tau^{est}_{H3K9} = \frac{\tau_{sim}}{8}$, where the total simulation time $\tau_{sim} \cong 1\ h$ was chosen to match the typical duration of embryonic stage 5 within NC14 (Fig.5c). In Fig. S5, we tested a perturbation that would slow down the speed of establishment ($\tau^{est}_{H3K9} = \frac{\tau_{sim}}{4}$).

**HP1 cluster detection & morphology analysis**

Phase-separated HP1 foci were identified through the computation of the local dimer occupancy ratio $\eta_i \in [0,1]$ at a given lattice site $i$, defined as

$$\eta_i = \frac{\sigma_i}{12} \sum_{j \in V(i)} \sigma_j.$$

In the above, $\eta_i = 0$ if site $i$ is not occupied by a HP1 dimer ($\sigma_i = 0$), and equals the fraction of neighboring sites $j \in V(i)$ occupied by other dimers if $\sigma_i = 1$. The discrete ensemble $P$ of sites delineating dense, phase-separated HP1 regions was specified by $P = \{\eta_i \geq 0.5\}$, although our results were found to be qualitatively insensitive to any choice of occupancy thresholds in the range $]\frac{1}{6}, 1[$. The topology of these domains was obtained from the adjacency list of sites $i \in P$ within the lattice, and the set of individual clusters $\{C_l\} = P$ was determined based on the connected components of $P$ via the NetworkX Python package (14).

The effective radius $R_l$ of a given cluster $C_l$ was defined as $R_l = \left(\frac{3N_l v_{site}}{4\pi}\right)^{\frac{1}{3}}$, with $N_l$ the number of HP1 dimers within the cluster, which corresponds to the radius of the virtual sphere with identical volume to $C_l$.
In the inset of Fig.4c, the mean droplet size $\langle R\rangle$ for Gao *et al.* (16) was extracted from the normalized fluorescence intensities $I$ reported in (16) via

$$\langle R(t)\rangle \sim \langle R\rangle_{max} \times I(t)^{1/3}.$$

In Fig.5f-h, for visual comparisons between predictions and experimental data from 4D microscopy, data and time were normalized by $\langle R\rangle_{max}$ and $\tau_{max}$ respectively such that $\langle R(t)\rangle = \langle R\rangle_{max}$ at $t = \tau_{max}$ with $R_{max}^{in\,vivo} \cong 1.7\,\mu\text{m}$, $R_{max}^{in\,silico} \cong 0.4\,\mu\text{m}$, $\tau_{max}^{in\,vivo} = 2160\,s$ and $\tau_{max}^{in\,silico} = 3000\,s$.
For a given cluster $C_l$, setting the cluster center of mass to the origin of the frame, the cluster gyration tensor $G_l$ was computed through

$$G_l^{mn} = \frac{1}{N_l}\sum_{i\in C_l} r_i^m r_i^n,$$

where $m, n \in \{x, y, z\}$ denote the projections of the coordinate vector $r_i$ of an arbitrary site $i$ onto the laboratory frame. Signifying by $\gamma_{1,2,3}$ the eigenvalues of $G_l$, the anisotropy parameter $A_l$ of the cluster reads as (15)

$$A_l = \frac{3}{2}\frac{\gamma_1^2+\gamma_2^2+\gamma_3^2}{(\gamma_1+\gamma_2+\gamma_3)^2} - \frac{1}{2},$$

which equals 0 in the case of spherically-symmetric foci and approaches 1 for needle-like, elongated clusters. The ensemble-averaged anisotropy $A_{sim}$ was finally obtained in the form

$$A_{sim} = \frac{1}{N_P}\sum_{C_l\in P} N_l A_l,$$

with $N_P = \sum_{C_l} N_l$ the total number of phase-separated HP1 dimers, and where the weights $N_l$ serve to subdue unphysically-large contributions from small foci.

The anisotropies obtained from 4D microscopy imaging data were similarly defined as

$$A_{exp} = \frac{1}{N_{vox}}\sum_{C_m} N_m(1 - S_m), \qquad \text{(S2)}$$

where the sum now runs over all distinct segmented foci $C_m$ (c.f. section "Live embryo imaging & analysis") and $N_{vox} = \sum_{C_m} N_m$. In Eq. (S2), $N_m$ denotes the number of voxels comprising an arbitrary focus $C_m$, whose sphericity $S_m$ may be related to the focus volume $V_m$ and surface area $\Sigma_m$ through

$$S_m = \frac{\pi^{1/3}(6V_m)^{2/3}}{\Sigma_m}.$$

With these conventions, Eq. (S2) ensures that $A_{exp} = 0$ for ideal spherical foci, while $A_{exp} \rightarrow 1$ indicates increasing deviations from sphericality – and thus establishes $A_{exp}$ as a direct experimental analog to $A_{sim}$. However, in Fig.5e, one may note that $A_{exp}$ has a different dynamical range than $A_{sim}$ (0.6-0.75 vs. 0-0.2, respectively).
The discrepancies between $A_{exp}$ and $A_{sim}$ and between $R_{max}^{in\,vivo}$ and $R_{max}^{in\,silico}$ could be due to optical artifacts from the limited microscopy resolution and/or to the intrinsic Rabl configuration of Drosophila chromosomes (11), which introduces an additional asymmetry in the genome 3D organization that is not accounted for in the simulations. Another potential source of disparity could be that our polymer model only explicitly describes the dynamics of a single long chromosome, out of the 6 present in live Drosophila embryos.

**Phase diagram & surface tension calculations**

The mean HP1 concentration $\rho_{HP1}^{dil}$ within the lattice region $D = \frac{L}{P}$ delimiting the dilute HP1 phase was defined as

$$\rho_{HP1}^{dil} = \frac{2\langle\sigma_i\rangle_D}{N_A v_{site}}, \qquad \text{(S3)}$$

in which $\langle\cdot\rangle_D$ denotes an ensemble average over all sites $i \in D$ lying outside of the phase-separated ensemble $P$. The surface tension $\gamma$ of the foci was then estimated based on the Bragg-Williams approximation of the lattice-gas partition function (17),

$$\gamma = \frac{k_B T N_A v_{site}}{4\delta^2}\left(\rho_{HP1}^{den} - \rho_{HP1}^{dil}\right) \times \left(\rho_{HP1}^{den} - \rho_{HP1}^{dil}\right),$$

where the concentration $\rho_{HP1}^{den}$ within HP1-dense regions was obtained similarly to Eq. (S3) based on the mean occupancy $\langle\sigma_i\rangle_P$ of sites within $P$. In practice, for all systems considered here, we find $\rho_{HP1}^{den} \cong \frac{2}{N_A v_{site}} \cong 600\ \mu M$. For the computation of such equilibrium quantities, simulations were first relaxed over $O(10^7)$ MC steps, and Eq. (S3) was further averaged over an additional $O(10^7)$ MC steps for each chosen set of parameters.

In all reported phase diagrams, each state point ($(\rho_{HP1}, J_{HP1-HP1})$ or $(\rho_{HP1}, J_{HP1-H3K9})$) corresponds to an independent simulation at steady state starting from a well-mixed system. The threshold concentration $\rho_{HP1}^0$ at the onset of droplet assembly was defined as the minimal concentration where the total fraction of HP1 present inside condensates is higher than 50 %, the remaining HP1 dimers in the background-phase allowing to compute $\rho_{HP1}^{dil}$ (Eq.S3). In the case of the pure HP1 system where the standard Flory-Huggins theory is valid, $\rho_{HP1}^{dil}$ could normally be assimilated to the low-density branch of the binodal, which corresponds to the curve in the $(\rho_{HP1}, J_{HP1-HP1})$ space delineating the region where condensates are a priori thermodynamically stable. The fact that $\rho_{HP1}^0$ is slightly higher than $\rho_{HP1}^{dil}$ suggests there may exist a small parameter range (between $\rho_{HP1}^{dil}$ and $\rho_{HP1}^0$) in our simulations where the system is metastable and remains well-mixed.

In the case of concentration-buffered systems, for which $\rho_{HP1}^{dil}$ is composition-independent, the equilibrium volume $V_{HP1}$ of HP1 foci in the phase-separated regime ($\rho_{HP1} > \rho_{HP1}^{dil}$) may be obtained via the lever rule for $\rho_{HP1}^{dil} \ll \rho_{HP1}^{den}$,

$$V_{HP1} = \frac{N N_A v_{site}^2}{2} \times \left(\rho_{HP1} - \rho_{HP1}^{dil}\right). \qquad \text{(S4)}$$

The dashed lines in Fig.3d represent the effective focus radii R associated with Eq. (S4) for both purified HP1α ($\rho_{HP1}^{dil} = \rho_{HP1\alpha}^{dil}$, lower red curve) and in the limit of infinite HP1 immiscibility ($\rho_{HP1}^{dil} \to 0$, upper red curve).

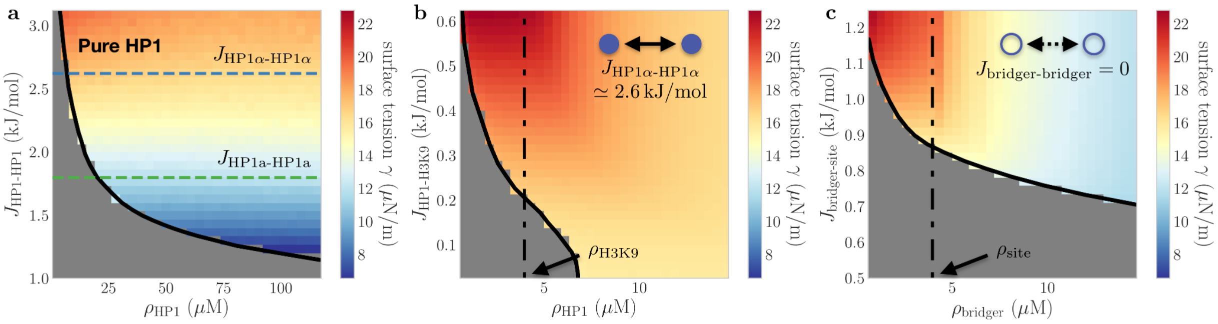


**Fig. S1: Surface tension of HP1 foci as predicted by the lattice-gas model.** (a) Equilibrium surface tension ($\gamma$) of HP1 foci as a function of HP1 density ($\rho_{HP1}$) and oligomeric affinity ($J_{HP1-HP1}$) computed for a purified HP1 system (see SI Methods). Note that $\gamma$ is largely independent of $\rho_{HP1}$ in this case and is further found to be comparable in magnitude to recent *in vivo* measurements for nuclear liquid condensates (20). (b) Effective surface tension of HP1-heterochromatin foci as a function of $\rho_{HP1}$ and HP1-H3K9 binding energy ($J_{HP1-H3K9}$) for HP1α ($J_{HP1-HP1} \cong 2.6 \frac{kJ}{mol}$) in the presence of mouse chromosome 19 (c.f. Fig.3). (c) Same as (b) in the case of non-interacting chromatin binding proteins ($J_{bridger-bridger} = 0$, c.f. Fig.S2).

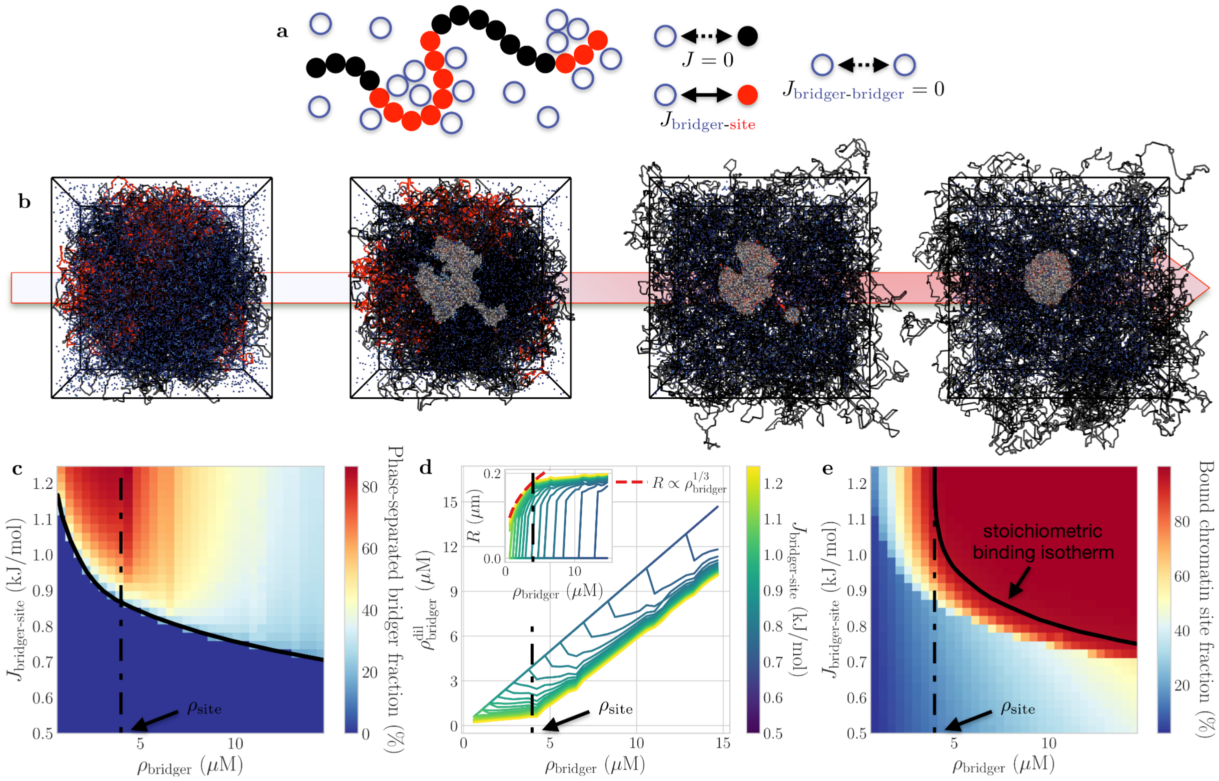


**Fig. S2: The "bridger" model.** (a) Sketch of the bridger-chromatin simulation scheme. Non-cooperative, diffusible "bridger" proteins do not self-interact ($J_{bridger-bridger} = 0$), but display an affinity for specific chromatin target sites with binding energy $J_{bridger-site}$, distributed along the chromosome as in Fig.1c. (b) Typical kinetic pathway of the simulations ($\rho_{bridger} \cong 8\ \mu M, J_{bridger-site} \cong 1 \frac{kJ}{mol}$). (c) Equilibrium phase diagram of the "bridger" model. Spontaneous phase separation occurs above a threshold concentration (black line), which is a function of $J_{bridger-site}$ (c.f. Fig.3b). (d) Dependence of the background (nucleoplasmic) bridger density $\rho^{dil}_{HP1}$ on the overall nuclear bridger concentration $\rho_{bridger}$ at various fixed $J_{bridger-site}$ (c.f. Fig.3c). Inset: Equilibrium radius of the corresponding foci. Dashed lines indicate the growth behavior expected of a concentration-buffered system based on the lever rule (c.f. Fig.3c). (e) is as in Fig.3e.

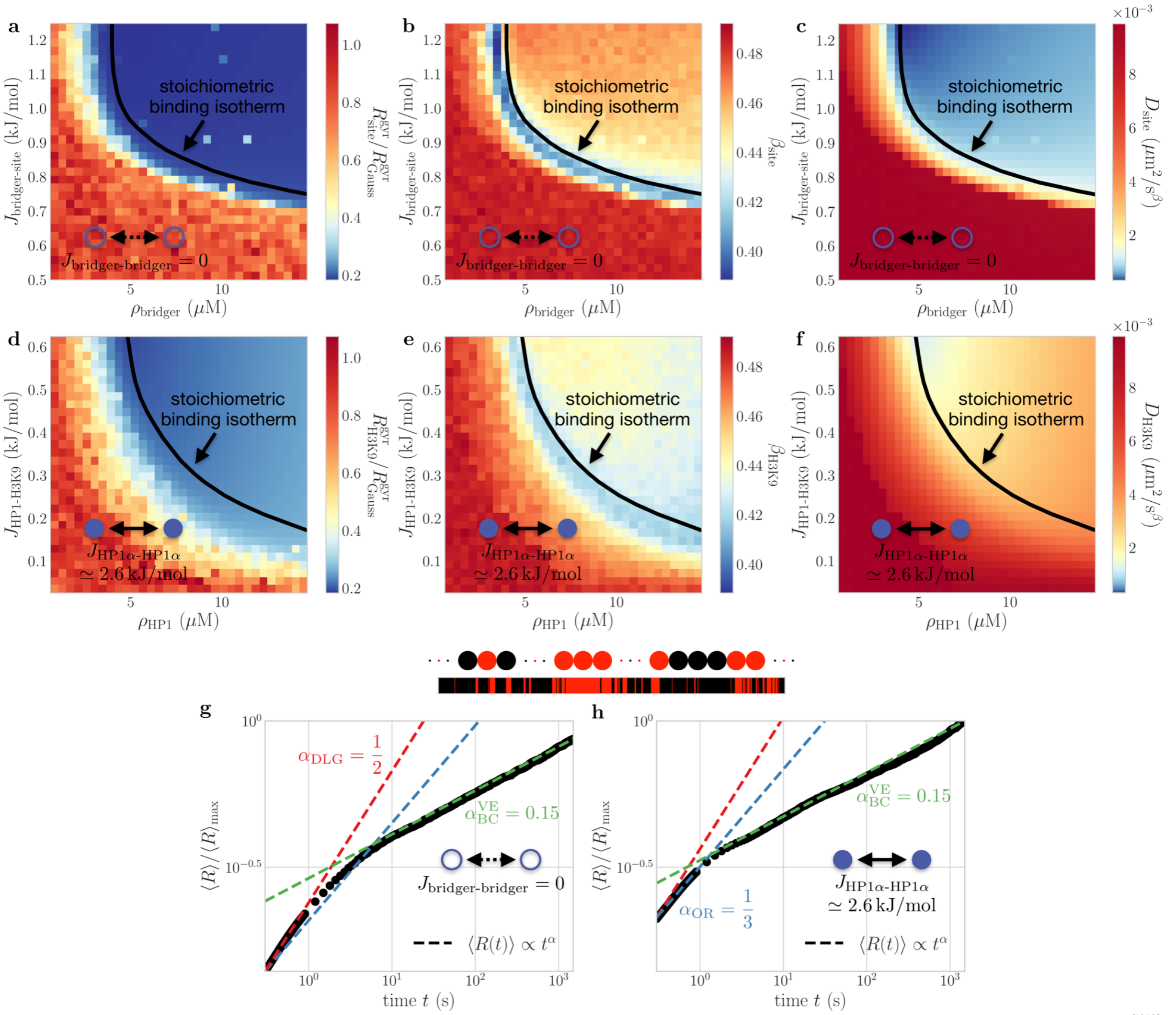


**Fig. S3: Comparison of the "bridger" and cooperative binding models.** (a) Normalized radius of gyration $R^{gyr}_{site}$ of the target chromatin domain as a function of $\rho_{bridger}$ and $J_{bridger-site}$ in the case of non-cooperative "bridger" proteins ($J_{bridger-bridger} = 0$, c.f. Fig.3f). (c) Chromatin diffusion exponent $\beta_{site}$ and (d) corresponding diffusion constant $D_{site}$ (c.f. Fig.3g). (d-f) are as in (a-c) for HP1α-like, self-interacting diffusible binders ($J_{HP1-HP1} \cong 2.6\ \frac{kJ}{mol}$). (a-f) Simulations were performed for the idealized telocentric mouse chromosome (Fig.1c). (g,h) Equilibration kinetics of the mean condensate radius $\langle R\rangle$ in the case of an idealized, human chromosome 19-like domain distribution. The data corresponding to the "bridger" (g) and cooperative binding models (h) are both normalized by the same maximal value $\langle R\rangle_{max} \cong 0.2\ \mu m$, as achieved by the self-interacting binders at the final timestep of the simulations. The initial growth period in the "bridger" model is ~3 times slower than the cooperative binding model. Both models then exhibit a coarsening dynamics driven by visco-elastic BC, but the corresponding coarsening prefactor of the "bridger" model is lower by ~15% than that of the cooperative binding model — consistent with the more strongly-inhibited diffusion kinetics exhibited by the encapsulated chromatin in this case (c.f. panels (c) and (f)).

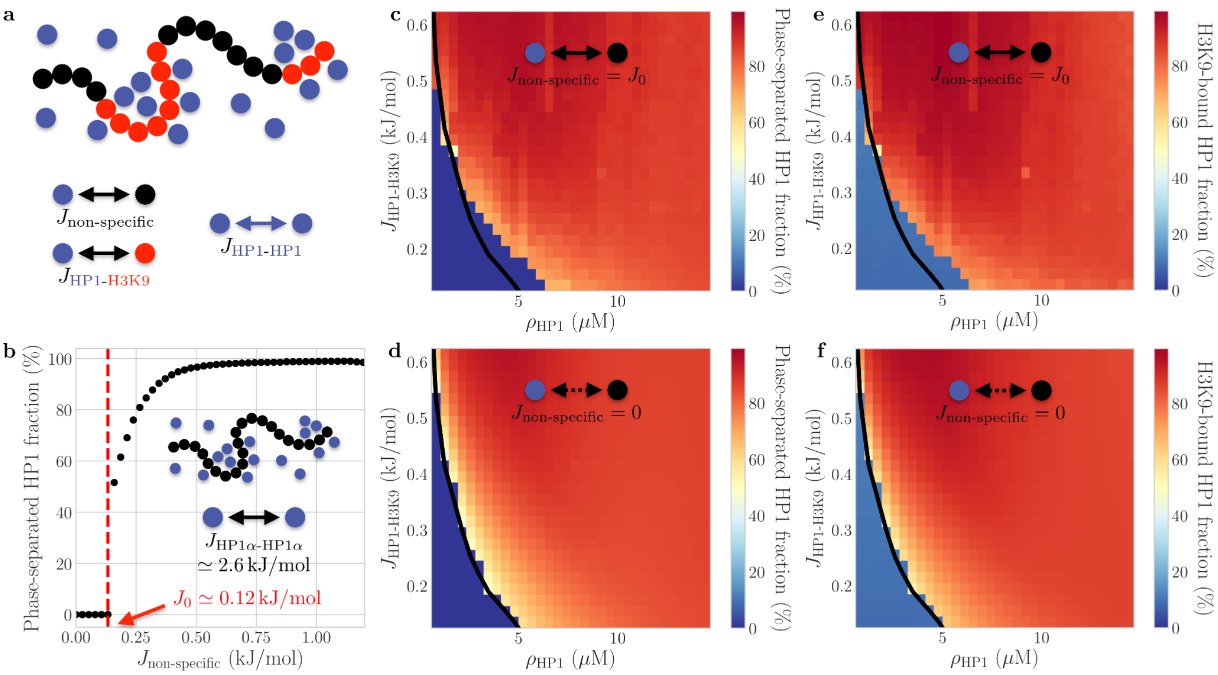


**Fig. S4: Role of non-specific interactions.** (a) Sketch of the cooperative binding model with non-specific interactions. HP1 dimers may now also bind non-H3K9me2/3 regions (black) with an arbitrary affinity $J_{non-specific}$. Other simulation details and parameters are as in Fig. 3 of the main text. (b) Non-specific interactions induce the condensation of HP1 and drive the associated collapse of a fully-unmethylated chromosome above a threshold value $J_0 \simeq 0.12 \frac{kJ}{mol}$ at physiological HP1α density ($\rho_{HP1} = 8\ \mu M$). $J_0$ can therefore be taken as an upper physical bound for $J_{non-specific}$, and the maximal potential effects of non-specific interactions in the context of the cooperative binding model may be probed by setting $J_{non-specific} = J_0$. (c,d) Phase diagram of the cooperative binding model with (c) and without (d) non-specific interactions, respectively. The black line denotes the approximate boundary of the two-phase region for $J_{non-specific} = 0$ in both cases in order to facilitate comparisons. Note that non-specific interactions slightly decrease the stability range of HP1-H3K9 condensates at low affinity $J_{HP1-H3K9}$, but otherwise have limited impact on the phase behavior. (e,f) The binding of HP1 onto H3K9 regions also correlates strongly with phase separation in both cases (c.f. (c,d)), indicating that HP1 condensation is driven chiefly by HP1-H3K9 specific interactions for $J_{non-specific} \leq J_0$.

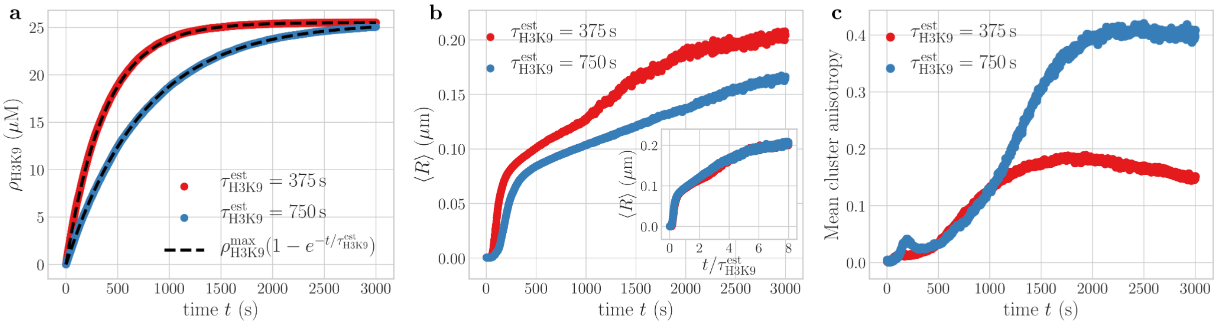


**Fig. S5: Role of H3K9 spreading kinetics on HP1 condensate growth.** (a) Two scenarios of H3K9me2/3 establishment with different uniform methylation rates: $\tau^{est}_{H3K9} = 375\ s$ (in red, c.f. Fig. 5c of the main text) and $\tau^{est}_{H3K9} = 750\ s$ (in blue). (b) The different methylation rates lead to qualitatively similar, but quantitatively distinct evolution kinetics of the mean radius $\langle R \rangle$ of HP1 foci. However, upon rescaling simulation times by $\tau^{est}_{H3K9}$, the two curves are found to closely collapse onto a unique master curve (inset). This indicates that the physics of condensate growth appears to be largely insensitive to the details of the H3K9 spreading process, consistent with a coarsening mechanism limited chiefly by the coupled diffusion kinetics of the HP1/polymer system (c.f. Fig. 5f of the main text). (c) The time evolution of the HP1 droplet anisotropy is also found to display similar trends in both scenarios, although slower H3K9 establishment kinetics are associated with a significantly larger loss of sphericity (blue curve). This may likely be attributed to the greater prevalence of late-stage focus nucleation events in this case, which promotes the formation of longer-ranged, persistent HP1 "bridges" (c.f. Fig. 5d of the main text).

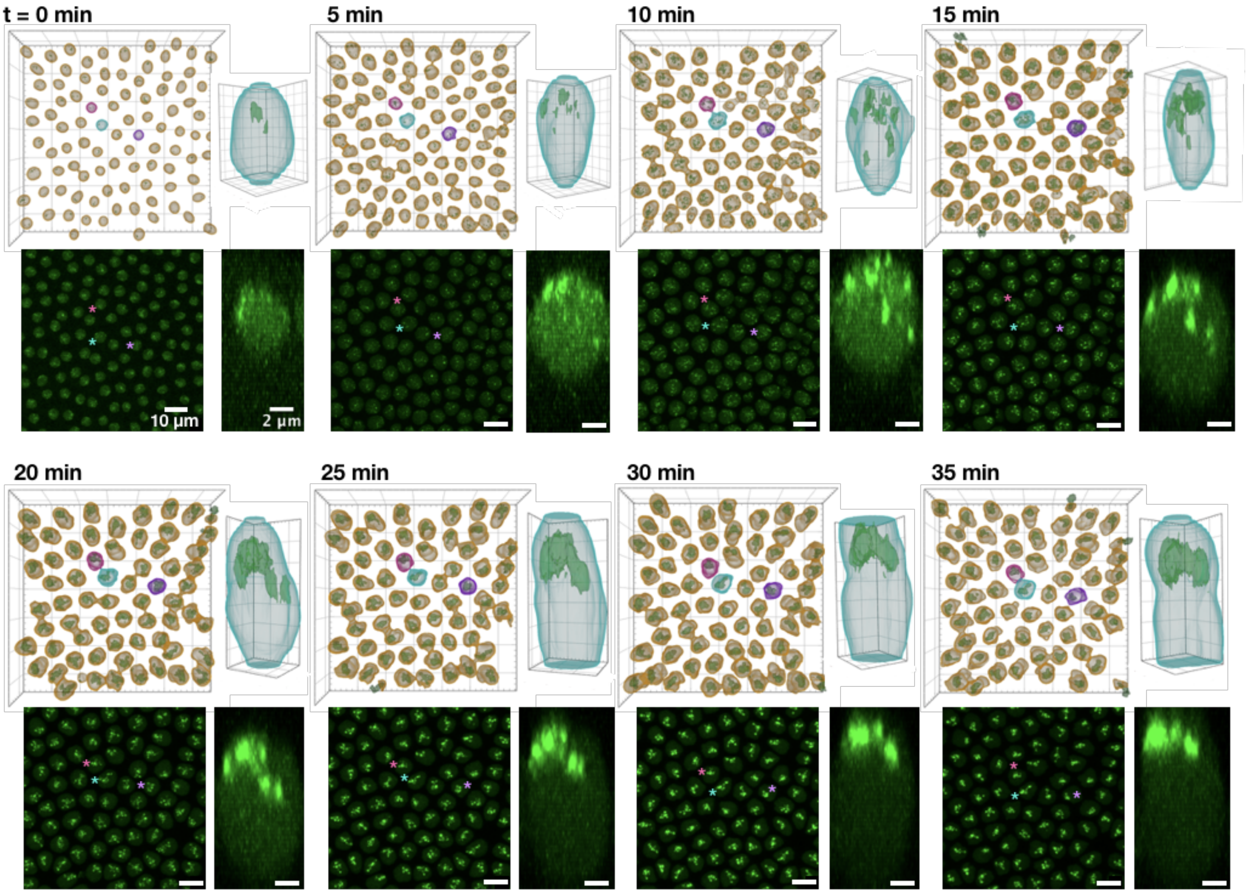


**Fig. S6: Time course images of nuclear cycle 14 drosophila embryos expressing GFP-HP1a.** (First and third rows) Arivis-generated 3D surface renderings of the whole field of view and a single nucleus. Full ROI grid is shown in 10 $\mu m$ increments across timepoints; individual nuclei renderings have a grid of 0.7 $\mu m$ ($t = 0$), 1 $\mu m$ ($t = 5\ min$), and 2 $\mu m$ ($t = 10{:}\ 35\ min$). (Second and fourth rows) Maximum intensity projections of GFP-HP1a in the whole field of view (z-projection, scale bar: 10 $\mu m$) and the corresponding single nucleus (axial-projection, scale bar: 2 $\mu m$).

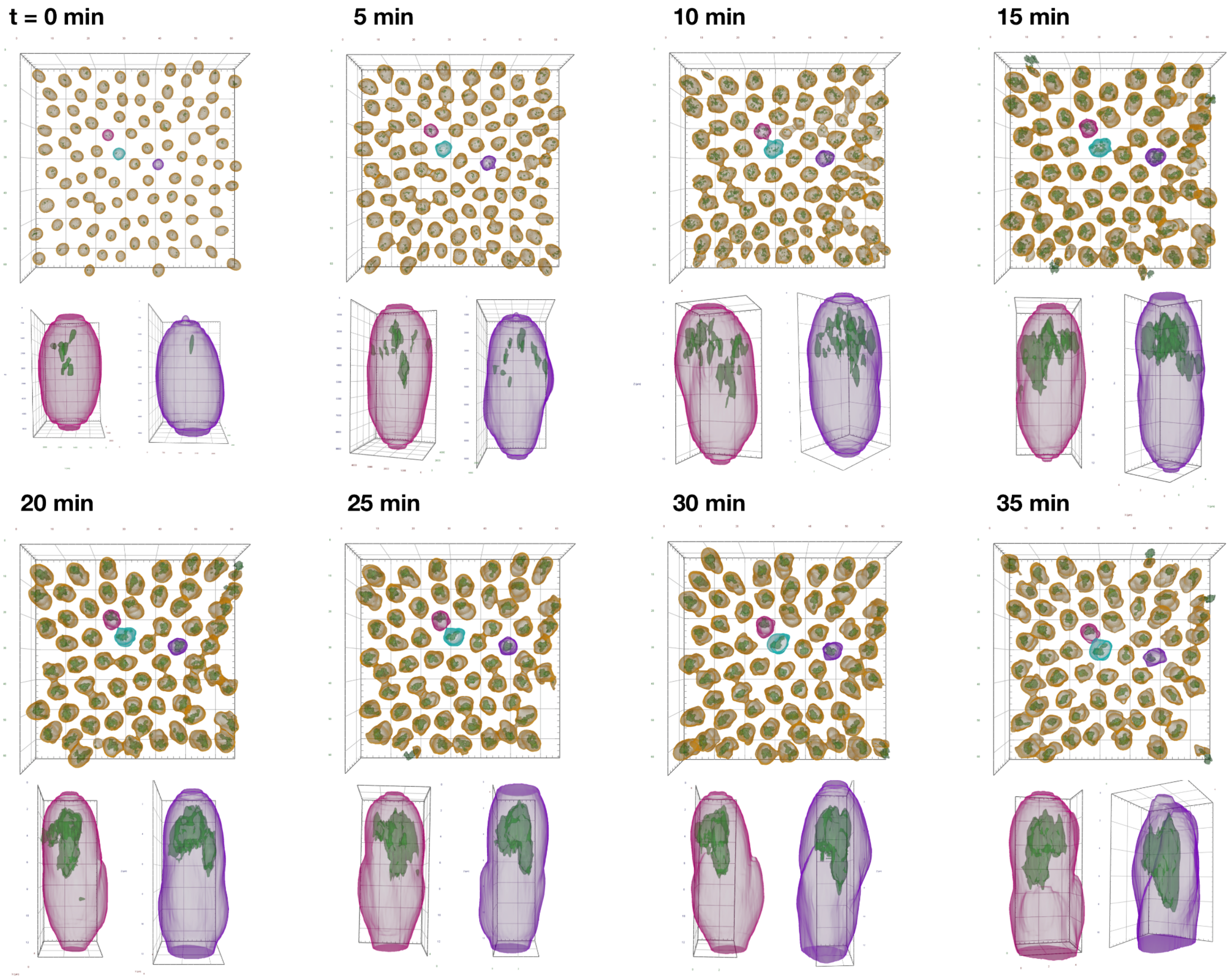


**Fig. S7: Time course images of Arivis-generated 3D surface renderings of the whole field of view and the indicated nuclei in nuclear cycle 14.** Full ROI grid is shown in 10 $\mu m$ increments across time points; individual nuclei renderings have a grid of 0.7 $\mu m$ ($t = 0$), 1 $\mu m$ ($t = 5\ min$), and 2 $\mu m$ ($t = 10{:}\,35\ min$).

**Movie S1 (separate file): Time-lapse video of the simulation associated with Fig. 2a.**

**Movie S2 (separate file): Time-lapse video of the simulation associated with Fig. 3a.**

**Movie S3 (separate file): Time-lapse video of the simulation associated with Fig. 4a.**

**Movie S4 (separate file): Time-lapse video of the simulation associated with Fig. 5b.**

### SI REFERENCES


1. S. K. Ghosh, D. Jost, How epigenome drives chromatin folding and dynamics, insights from efficient coarse-grained models of chromosomes. *PLoS Comput. Biol.* **14**, e1006159 (2018).

2. F. Erdel, *et al.*, Mouse Heterochromatin Adopts Digital Compaction States without Showing Hallmarks of HP1-Driven Liquid-Liquid Phase Separation. *Mol. Cell* **78**, 236–249.e7 (2020).

3. S. Sati, *et al.*, 4D Genome Rewiring during Oncogene-Induced and Replicative Senescence. *Mol. Cell* **78**, 522–538.e9 (2020).

4. M. J. D. Powell, An Efficient Method for Finding the Minimum of a Function of Several Variables Without Calculating Derivatives. *Comput. J.* **7**, 155–162 (1964).

5. M. M. C. Tortora, H. Salari, D. Jost, Chromosome dynamics during interphase: a biophysical perspective. *Curr. Opin. Genet. Dev.* **61**, 37–43 (2020).

6. L. Schmiedeberg, K. Weisshart, S. Diekmann, G. Meyer Zu Hoerste, P. Hemmerich, High- and low-mobility populations of HP1 in heterochromatin of mammalian cells. *Mol. Biol. Cell* **15**, 2819–2833 (2004).

7. K. Müller-Ott, *et al.*, Specificity, propagation, and memory of pericentric heterochromatin. *Mol. Syst. Biol.* **10**, 746 (2014).

8. S. Alberti, A. Gladfelter, T. Mittag, Considerations and Challenges in Studying Liquid-Liquid Phase Separation and Biomolecular Condensates. *Cell* **176**, 419–434 (2019).

9. A. R. Strom, *et al.*, Phase separation drives heterochromatin domain formation. *Nature* **547**, 241–245 (2017).

10. J. Bonnet, *et al.*, Quantification of Proteins and Histone Marks in Drosophila Embryos Reveals Stoichiometric Relationships Impacting Chromatin Regulation. *Dev. Cell* **51**, 632–644.e6 (2019).

11. V. E. Foe, B. M. Alberts, Reversible chromosome condensation induced in Drosophila embryos by anoxia: visualization of interphase nuclear organization. *J. Cell Biol.* **100**, 1623–1636 (1985).

12. D. V. Fyodorov, Acf1 confers unique activities to ACF/CHRAC and promotes the formation rather than disruption of chromatin in vivo. *Genes Dev.* **18**, 170–183 (2004).

13. K. H.-C. Wei, C. Chan, D. Bachtrog, Establishment of H3K9me3-dependent heterochromatin during embryogenesis in Drosophila miranda. *eLife* **10** (2021).

14. A. Hagberg, P. Swart, D. Schult, Exploring network structure, dynamics, and function using NetworkX. *Proceedings of the 7th Python in Science Conference*, 11–15 (2008).

15. D. N. Theodorou, U. W. Suter, Shape of unperturbed linear polymers: polypropylene. *Macromolecules* **18**, 1206–1214 (1985).

16. Y. Gao, M. Han, S. Shang, H. Wang, L. S. Qi, Interrogation of the dynamic properties of higher-

order heterochromatin using CRISPR-dCas9. *Mol. Cell* **81**, 4287–4299.e5 (2021).

17. G. W. Woodbury, General Equation for the Surface Tension of the Lattice Gas. *J. Chem. Phys.* **51**, 1231–1235 (1969).

18. A.V. Probst, E. Dunleavy, G. Almouzni, Epigenetic inheritance during the cell cycle. *Nat Rev Mol Cell Biol* **10**, 192–206 (2009).

19. C. Alabert, *et al*, Two distinct modes for propagation of histone PTMs across the cell cycle. *Genes Dev.* **29**, 585–590 (2015).

20. C. M. Caragine, S. C. Haley, A. Zidovska, Surface Fluctuations and Coalescence of Nucleolar Droplets in the Human Cell Nucleus. *Phys. Rev. Lett.* **121**, 148101 (2018).
